# 2D spatial tissue analysis often misrepresents true biological patterns of 3D tissues

**DOI:** 10.64898/2026.01.21.700966

**Authors:** David Le, David I Kaplan, Anna S Trigos

## Abstract

Spatial omics technologies and analysis methods that profile tissue sections are widespread in biomedical research, but how well these methods capture 3D tissue biology is unknown. Using 3D spatial simulations and data, coupled with virtual tissue sectioning, we found that spurious associations, misrepresentation of biological trends and false negative findings are common in 2D spatial analyses. Understanding the limitations of 2D spatial analyses is needed to avoid misinterpreting research findings.

## Main text

We aimed to investigate potential discrepancies between spatial analysis results obtained from 3D samples and 2D tissue sections. We developed SPIAT-3D, an R toolkit that enables the use of the third dimension in the calculation of 14 commonly used 2D metrics,^1–7^ including average minimum distance (AMD), mixing score (MS), normalised mixing score (NMS), average cells in neighbourhood proportions (ACIN), average neighbourhood counts (ANC), average neighbourhood entropy (ANE), cross-K function (CK), cross-L function (CL), cross-G function (CG), co-occurrence (COO), proportion-based spatial autocorrelation (PBSAC), proportion-based prevalence (PBP), entropy-based spatial autocorrelation (EBSAC), entropy-based prevalence (EBP) (Supp Note 1, Supp Fig. 1). We compiled four publicly available 3D spatial samples: a human colorectal cancer sample generated with CyCIF,^8^ a human metastatic lymph node sample generated with Open-ST,^9^ and a mouse brain cortex and hypothalamus sample generated with MERFISH.^10^ We performed *in silico* tissue sectioning (i.e. virtual sectioning) (Fig. 1a) to obtain 2D sections of each of the 3D samples, replicating what occurs experimentally, and calculated all 14 spatial metrics on the 3D samples and matching 2D sections (Fig. 1b, Supp. Fig 2). The 2D results varied greatly depending on the selected slice, resulting in a wide range of percentage differences between 2D results and the reference 3D values (between –416.21% and 363.61% across datasets and metrics, Table S1), and some metrics could either over or underestimate the 3D values depending on the selected slice.

**Figure 1.**
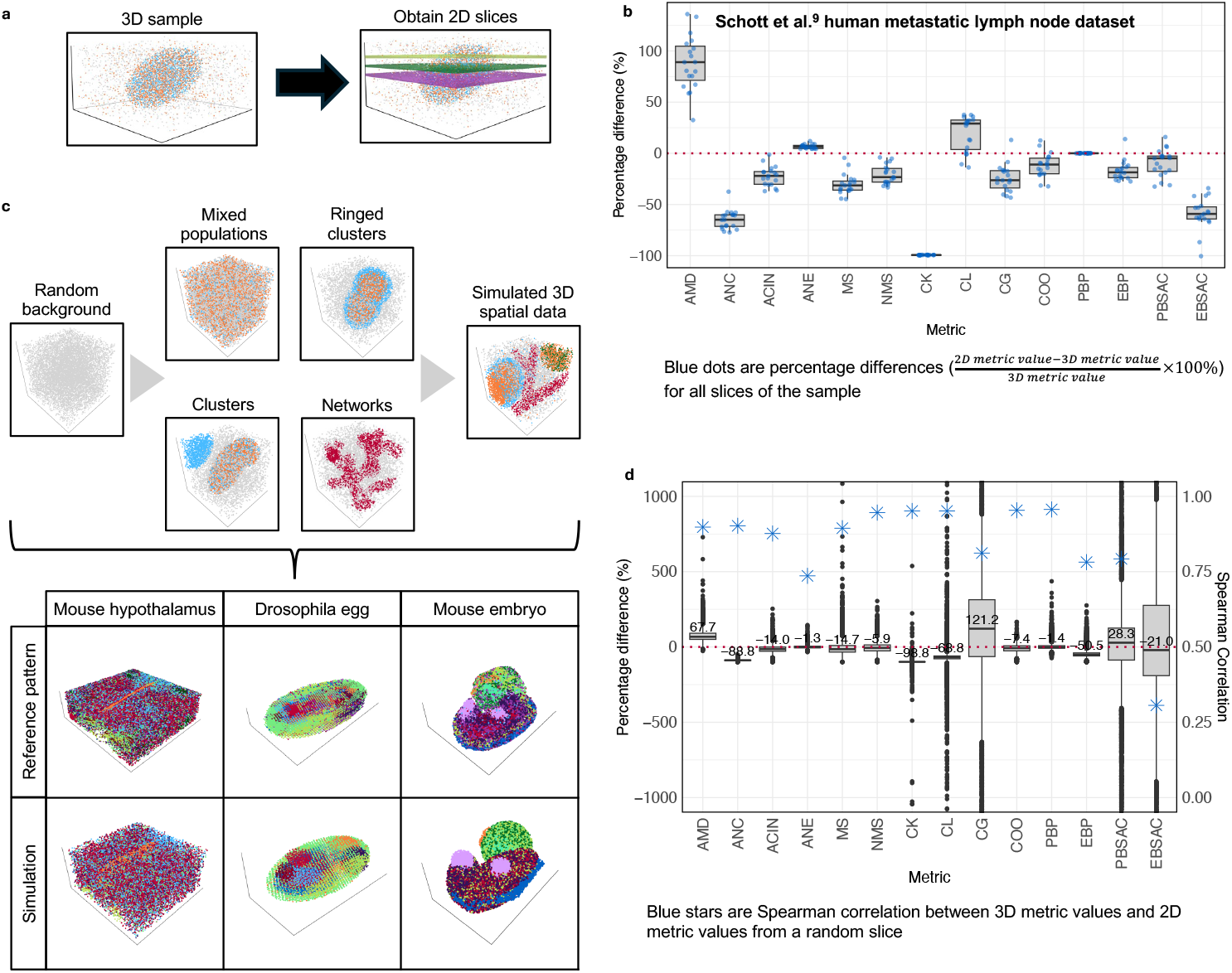
Comparison of spatial analysis in matched 3D samples and 2D sections. **(a)** Virtual sectioning of 3D simulations to generate 2D tissue sections. **(b)** Percentage difference of the 2D metric values for each section compared to their 3D counterparts. **(c)** spaSim-3D. Components used to build a 3D spatial simulation (top). spaSim-3D can simulate complex 3D structures composed of cells of defined categorical types (bottom). **(d)** Percentage difference of the 2D metric values from 10,000 3D simulations and a randomly selected, matched 2D section.

We hypothesized that the discrepancy between 2D and 3D values might be linked to tissue complexity and spatial properties of the sample. To investigate, we developed spaSim-3D, a simulator of spatial tissue patterns in 3D (Fig. 1c, Supp. Note 2). First, we generated a cohort of 10,000 simulations by randomly varying all parameter combinations (Simulated Dataset One, Methods section 3.1), followed by virtual sectioning and calculation of the spatial metrics (Methods section 3.2, Supp. Fig. 3). The Spearman correlation between 2D and 3D values across all samples was above 0.85 in 9/14 metrics (Supp. Fig. 4, Table S2), and this increased to 13/14 when the average across multiple 2D sections were taken (Table S2). However, correlations tended to be higher in simulations with simpler spatial structures (e.g. ellipsoids) than in simulations with network and ringed structures (Table S2). Despite high correlations, the percentage difference ranged from -22,092,405% to 8,544,782% and showed similar trends to those obtained from the biological data (Fig. 1b and 1d, Table S3). Averaging across 2D sections reduced the variability of 2D results in 11/14 metrics, although the percentage difference still remained high (Supp. Fig. 5a, Table S3). Concerningly, the percentage difference of the 2D scores was not constant across the spectrum of 3D values, with often complex associations of higher or lower error depending on the raw magnitude of the 3D values (Supp Fig. 6). To further understand how the spatial composition affected the 2D vs 3D relationship, we generated 24,000 simulations across six tissue structure configurations and controlling seven individual parameters (Simulated Dataset Two, Methods section 3.1). As before, we performed virtual sectioning and calculated the spatial metrics in both the 3D and 2D samples. The associations between 3D and 2D spatial metrics were sensitive to the spatial properties of the samples (Supp. Fig. 7), including the volume of structures, the distance separating structures, the width of network edges, the proportions of cell types in the background, the level of cell mixing and the presence of ringed structures. However, whether or how these properties affected the 2D spatial results depended on the specific spatial metrics (Supp Fig. 7).

Overall, we identified that 2D spatial analysis results did not have a simple relationship with 3D results, and could be often considered spurious. The diversity of spatial arrangements, tissue structures and cell distributions resulted in complex associations between 3D and 2D results, which were unique to each spatial metric. Common strategies such as averaging across tissue sections of a single sample or increasing sample sizes did not significantly overcome these limitations.

Next, we investigated how the spurious nature of the 2D metrics could affect the results of analyses comparing sample groups. We created four collections of simulations (Simulated Dataset 3), each with 200 sets of 3D simulations, where each set was composed of two groups, for a total of 32,000 simulations (Fig. 2a). Collections were designed to test either the false positive (FPR) or false negative rate (FNR) (Fig. 2b, left) of 2D analyses by comparing the results obtained with 3D data. For the FPR collections (FPRc), both sample groups in each pair were simulated using similar parameter ranges (Methods section 5). In contrast, for the FNR collections (FNRc), the groups of each pair were simulated with distinct spatial parameter values. As before, we performed virtual sectioning and randomly selected a 2D section for each 3D simulation. The FPR for both the 3D and 2D datasets was ∼ 0.05 across all spatial metrics tested, regardless of tissue complexity (Fig. 2b), indicating no evidence of an increase in spurious discoveries in 2D compared to 3D analyses in data. However, we identified that significant differences between groups in the FNRc were more likely to be found when using the 3D data compared to the 2D data, and this effect was exacerbated when simulations of more complex tissue structures, such as networks, were used (Figure 2b).

**Figure 2.**
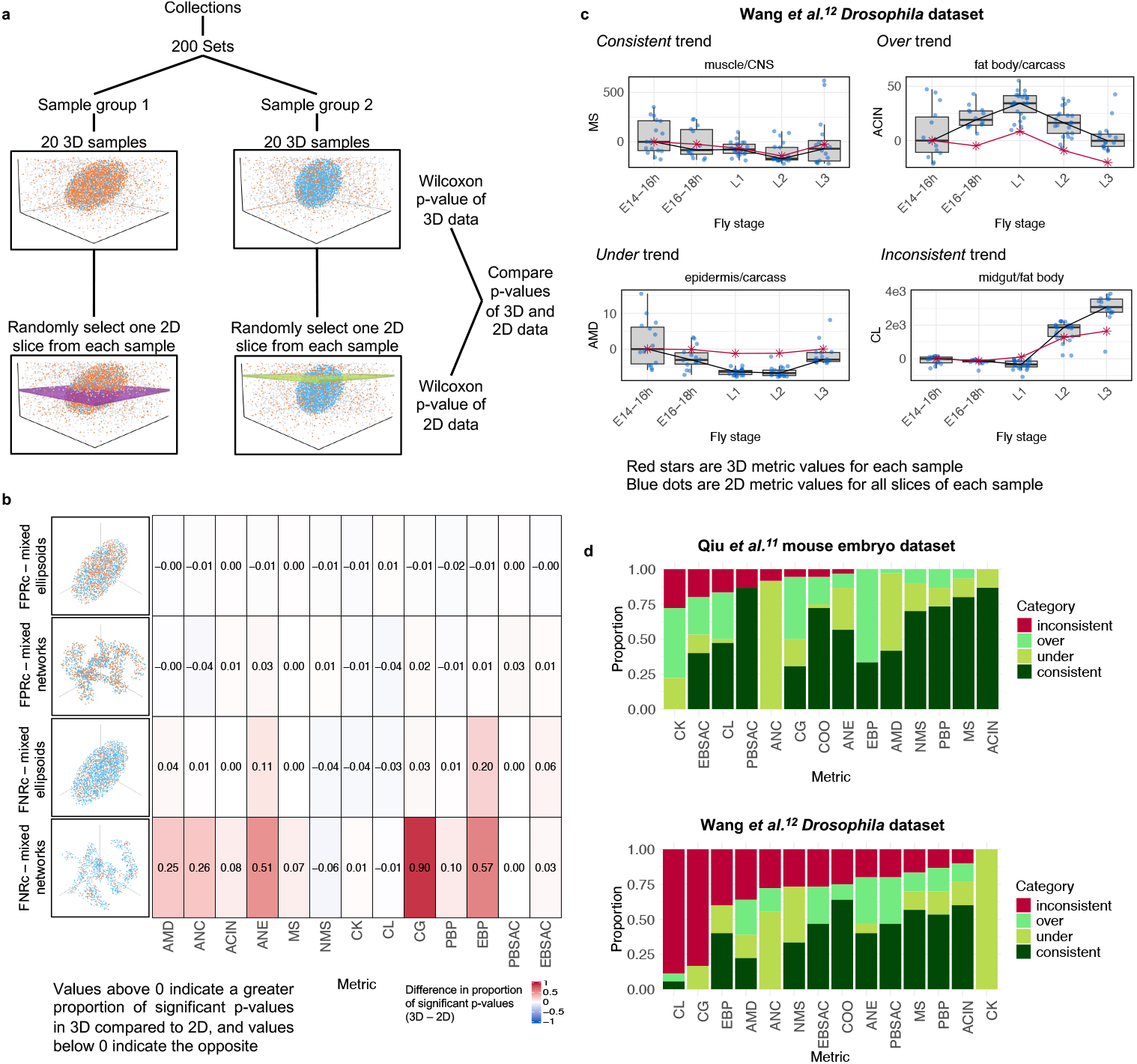
Analysis of serial 2D sample groups limits the identification of trends and leads to spurious findings. **(a)** Collections, sets and sample groups. **(b)** Difference between the proportion of significant p-values (below 0.05) obtained with 3D and 2D metrics. **(c)** Examples of categorical trends. **(d)** Proportion of the incidence of consistent, over and underestimating, and inconsistent trends of 2D metrics compared to 3D metrics across all pairs of cell types.

**Figure 3.**
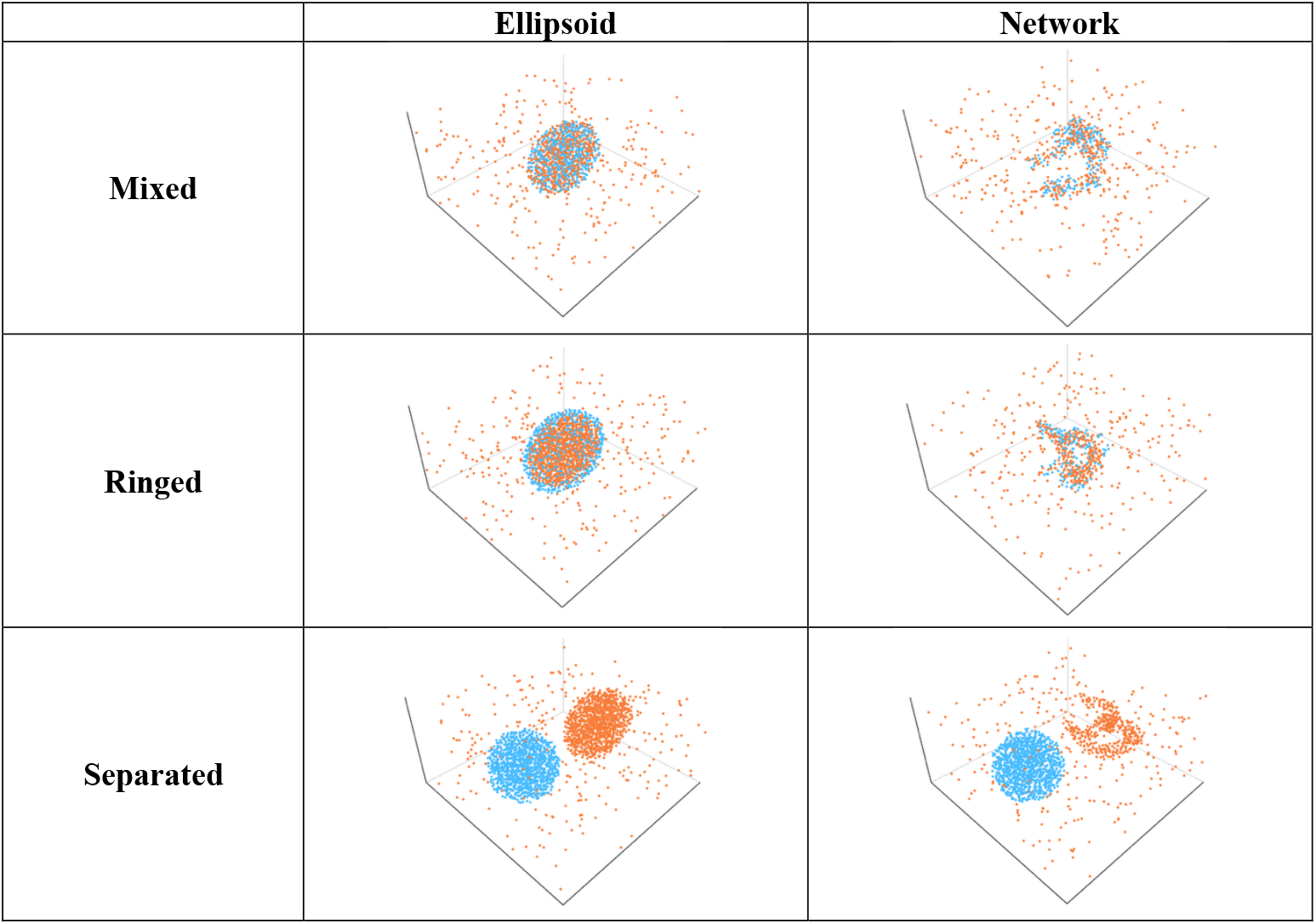
Example simulations for each categorical parameter set. Cell types: A (orange), B (blue) and O (predominant background cell type not shown for clarity). A mixed cluster arrangement is a cluster with both cell types A and B. A ringed cluster arrangement is a cell type A cluster surrounded by a cell type B ring. Separated cluster arrangement are two distinct clusters, cluster1 comprised of only cell type A, and cluster2 comprised of only cell type B (fixed sphere). Cluster shapes include ellipsoid and network.

Finally, to determine how analysis of time series data would be affected by performing analyses in 2D, we used a mouse dataset from Qiu et al.^11^ generated using spateo of two mouse embryo samples 9.5 hours and 11 hours after birth, and a dataset from Wang et al.^12^, who used Stereo-seq to profile 5 consecutive stages of *Drosophila* development. We determined whether the 2D spatial metric scores were either *consistent* (within a similar range) with the 3D scores, were consistently *over-* or *under*estimating the 3D values, or if the 3D and 2D values showed *inconsistent* trends (Fig 2c, Methods section 6.2). Although a subset of metrics revealed trends in the 2D analyses that were generally consistent with those identified from 3D samples (COO, MS, PBP, ACIN), metrics with the highest incidence of inconsistent trends differed between datasets (Fig 2d). These results point towards a limited reliability of 2D spatial metrics in identifying trends between multi-sample experiments Our work showcases strategies to test the behaviour of spatial methods when they are applied on 2D data compared to 3D. We demonstrate inherent limitations and confounders of 2D spatial analyses, which are critical to understand before ample interpretation of findings or consideration of clinical applications.

## Methods

### 1. Spatial analysis in 3D with SPIAT-3D

SPIAT-3D is a comprehensive analytical toolkit for computing spatial patterns in 3D tissue structures. We leveraged and expanded existing metrics present in SPIAT and adapted them to effectively handle 3D spatial data.^1^ The introduction of a third dimension necessitated innovative adaptations to these metrics, often involving the incorporation of the z-coordinate into metric calculations. Additionally, we introduced several new metrics exclusive to SPIAT-3D to address the unique challenges and opportunities presented by 3D data. These new metrics include neighbourhood counts, neighbourhood entropy and co-occurrence.^6^ A detailed description of all SPIAT-3D algorithms can be found in Supplementary Note 1.

The SPIAT-3D cell colocalization metrics used for analysis of 3D tissue data were the following: minimum distance between cells, neighbourhood counts, cells in neighbourhood, neighbourhood entropy, mixing score (MS), normalised mixing score (NMS), cross K-function (CK), cross L-function (CL), cross G-function (CG) and co-occurrence (COO). For our required downstream analysis, a single value from each metric was required for each 3D tissue sample. Therefore, for cell colocalization metrics which provided an output value for each cell in the 3D tissue sample (minimum distance between cells, neighbourhood counts, cells in neighbourhood and neighbourhood entropy), we calculated an average for those metrics across each cell, resulting in the average minimum distance between cells (AMD), average neighbourhood counts (ANC), average cells in neighbourhood (ACIN) and average neighbourhood entropy (ANE), respectively.

Metrics which used an input gradient of radius values (ANC, ACIN, ANE, MS, NMS, CK, CL, CG, COO) were applied to each 3D tissue sample for a gradient of radius values from 20 to 100 with a step size of 10. We selected this radius range as we found radii below 20 could not consistently capture any interactions between cells, whilst radii above 100 showed little additional variation. We selected a step size of 10 after observing that finer increments (e.g., 1 – 5) produced highly similar trends. Then, the area under the curve was calculated across the gradient of radius values using the Trapezoidal Rule.

The SPIAT-3D spatial heterogeneity metrics used for 3D spatial analysis were prevalence and spatial autocorrelation. These metrics first require each 3D tissue sample to be divided into a grid of rectangular prisms. For all analyses, we used 10 splits along each axis (x, y and z), producing a 10 by 10 by 10 grid and therefore 1,000 rectangular prisms per 3D sample. We selected 10 splits because this resolution consistently captured the spatial trends expected from each spatial heterogeneity metric and increasing the number of splits did not meaningfully change the results. Then, the proportion value or entropy value was calculated in each rectangular prism. This resulted in 4 distinct spatial heterogeneity metrics: proportion-based prevalence (PBP), proportion-based spatial autocorrelation (PBSAC), entropy-based prevalence (EBP), entropy-based spatial autocorrelation (EBSAC). PBP and EBP are implemented in SPIAT-3D to use an input gradient of threshold values from 0.01 to 1 with a step size of 0.01. This was converted to a single value for downstream analysis by calculating the area under the curve across the gradient of threshold values using the Trapezoidal Rule.

### 2. 3D spatial data simulator – spaSim-3D

spaSim-3D is a spatial simulator that can generate a variety of 3D tissue structures, using a similar step-by-step process as those implemented in spaSim.^1^ We used spaSim as a foundation and introduced significant modifications to accommodate 3D simulations. This included incorporating the z-coordinate of cells, and adapting shape generation techniques from 2D to 3D to develop novel 3D simulation algorithms.

As the parameters used to generate simulations can be largely determined by the user, spaSim-3D facilitates highly customisable tissue structure simulations. This allows spaSim-3D to generate different spatial structures that are commonly found in tissues. Furthermore, the user can mix and match different 3D simulation functions to simulate intricate tumour tissue structures. Detailed descriptions of the methods used to generate these simulations are described in Supplementary Note 2.

### 3. Creation of 3D simulation images, virtual slicing and comparison of matched 3D and 2D simulated data

Here we aimed to determine whether 2D analysis can adequately represent 3D data. To achieve this, we generated thousands of 3D simulations with varying parameters using spaSim-3D and applied SPIAT-3D metrics to these simulations. We then used virtual slicing to select multiple 2D slices from each 3D simulation and calculated the corresponding 2D metrics. These were then compared to understand the behaviour of factors such as the specific metric used and any association with underlying patterns within the tissue data.

#### 3.1. 3D simulation generation

In this section, we generated two datasets of 3D tissue simulations. Simulations in both datasets were composed of different types of cell clusters made up of two generic cell types: cell type A and cell type B. The background of all simulations consisted of another cell type: cell type O.

In Simulated Dataset One, simulations were designed to mimic the complexity and variability of real tissue biology. To do this, we varied most spaSim-3D simulation parameters. Fixed parameters, such as the dimensions of the simulation space and the total number of cells remained the same for all simulations (Table S4). However, categorical parameters, were randomly sampled, and continuous parameters were uniformly sampled from pre-defined ranges simultaneously (Table S5, Table S6). In total, Simulated Dataset One consisted of 10,000 simulations with a wide range of parameter combinations.

In Simulated Dataset Two, simulations were generated such that only one parameter was being changed at a time, to allow determining the effect of specific parameters on the 2D-3D relationship. Similar to Simulated Dataset One, fixed parameters remained the same for all simulations (Table S4). Categorical parameters (the shape and arrangement of clusters) were tested by creating 6 categorical parameter sets: mixed-ellipsoid, mixed-network, ringed-ellipsoid, ringed-network, separated-ellipsoid, and separated-network (Table S5, Figure 3). Next, for each categorical parameter set, we created 1000 simulations by uniformly sampling a value for one continuous parameter at a time, while keeping all other parameters fixed (Table S6). Each categorical parameter set had 4 continuous parameters that could be varied (Table S7), generating in total 24,000 simulations for Simulated Dataset Two.

#### 3.2. Virtual 2D slicing process

Next, to obtain 2D slices from each 3D simulation, we extracted orthogonal slices along the x-y plane. Each slice had a width of 10 units, corresponding to the ‘minimum distance between cells’ parameter (Table S4) used when generating the background cells of each simulation, ensuring that each slice contained, roughly, a single layer of cells. This approach resulted in approximately 1,000 cells per slice. For Simulated Dataset One, we extracted 13 consecutive slices from z = 85 to z = 215 and randomly selected one 2D slice for each 3D simulation, to provide a one-to-one comparison between 2D and 3D metrics on biologically realistic spatial simulations. This process mimics what is routinely done when generating spatial –omics data where a single section is used for profiling. For Simulated Dataset Two, we extracted three 2D slices to investigate the impact of slice position on 2D spatial metrics. The first slice was positioned directly through the centre of the simulation (z = 145 to z = 155), while the second and third slices were located 30 and 60 units above the first, respectively (z = 175 to z = 185 and z = 205 to z = 215).

To also test the impact of slice position in Simulated Dataset One, we sampled 3 slices from the 13 consecutive slices with the same z-coordinates as the 3 slices extracted for Simulated Dataset Two.

#### 3.3. 3D and 2D spatial metric comparison

Finally, to compare spatial metrics derived from 3D and 2D analyses, we applied SPIAT-3D metrics (see specified parameters in Table S8) to each 3D simulation in Simulated Dataset One and Simulated Dataset Two. As input to each metric, we used cell types A and B of the simulations. For metrics where a reference cell type and target cell type was needed, such as AMD, CK, PBSAC, we set cell type A as the reference cell type and cell type B as the target cell type. For other metrics which require a set of cell types as input, such as EBSAC and EBP, we used both cell type A and B. For each 2D slice, we applied the corresponding SPIAT metrics (or derived equivalent 2D metrics where necessary). The results from both 3D and 2D analyses were then compared to assess the accuracy of 2D representations of 3D tissues. For Simulated Dataset One, on top of analysing a randomly selected 2D slice and 3 2D slices at specific positions, all 13 consecutive slices were also analysed in 2D and an average was obtained to determine if averaging the 2D metric values calculated from the slices can improve the 2D-3D metric relationship.

### 4. Comparing spatial metrics in 3D and 2D publicly available data

To compare 2D and 3D spatial analysis in real biological data, we applied SPIAT-3D metrics to three publicly available 3D datasets.

#### 4.1. Publicly available data acquisition and preprocessing

Three public 3D tissue datasets were used, as described below. In each of these datasets, the 3D data was initially acquired as 2D sections, which could then be stacked to form 3D spatial datasets. For each dataset, the x, y, z coordinates and cell type identity of every cell from each 2D section was extracted and combined to construct three comprehensive 3D datasets. All datasets already included categorical cell type labels for each cell, which we used for our analysis as these are the required input to SPIAT and SPIAT-3D.

##### 4.1.1. Human colorectal cancer 3D dataset

We used a publicly available colorectal cancer dataset generated by Lin et al.^8^ In total, 93 colorectal cancer samples were collected and analysed in this study, but 3D tissue reconstruction was performed on only the first tissue sample, which we used in our study. For this sample, 106 serial sections were cut and 22 H&E and 25 CyCIF images were collected (other sections were skipped). In our analyses, we used the data from the 25 CyCIF-stained sections. Here, for each section, a separate CSV file was available containing the x, y coordinates and cell identity of every cell. Each file had been labelled with its serial section number to distinguish between sections. To obtain the z-coordinate of each cell, we used the serial section number and slice thickness provided.^8^ This allowed us to calculate the z-coordinate for each serial section which was assigned to the corresponding cells found in each section. After adding the z-coordinate to each CSV file, all CSV files were merged into a singular file containing the x, y, z coordinates and cell identity of every cell. In total, the final 3D dataset contains 12,697,850 cells.

##### 4.1.2. Human metastatic lymph node 3D dataset

We used a 3D dataset from Schott et al.,^9^ who used Open-ST, to analyse a 3D human metastatic lymph node tissue. In this work, the authors extracted 36 sections from the human metastatic lymph node tissue, of which 19 sections were profiled using Open-ST, 11 were stained with H&E, 1 was profiled with immunofluorescence and 5 sections remained for validation.

For our study, we accessed the publicly available H5AD file, which contained the x, y, z coordinates and cell type identity of every cell from the 19 Open-ST profiled sections. In total, the dataset had 1,097,769 cells.

##### 4.1.3. Mouse brain cortex 3D dataset and mouse hypothalamus 3D dataset

We obtained mouse brain tissue data from Fang et al.,^10^ generated using MERFISH combined with 3D confocal microscopy to examine the brain cortex and hypothalamus. This spatial-omics approach provides cell coordinates and transcriptomic profiles without the need for serial sectioning. For our analysis, which required both 3D tissue samples and matched 2D representations, we extracted the x, y, z coordinates and cell type identities of each cell from the provided CSV files. To generate 2D data, we discretised the z-axis into 10 µm bins: cells with z-coordinates between 0–10 µm were assigned to 0 µm, those between 10–20 µm to 10 µm, and so forth. For the mouse brain cortex sample, 10 bins or sections were extracted, while for the mouse hypothalamus sample, 20 sections were extracted, as this section was thicker. 35,149 cells were contained in the mouse cortex sample while 79,589 cells were present in the mouse hypothalamus sample.

##### 4.1.4. 3D and 2D spatial analysis of public data

We applied SPIAT-3D metrics with the same parameters used in Table S8 (apart from the cell type inputs) to the each publicly available 3D dataset. The cell types available in each dataset was used as input for each metric.

For the 3D human colorectal cancer dataset, we used all 21 unique cell type identity annotations provided by Lin et al.,^8^ providing 441 cell pairs (from 21^2^) as input for the spatial metrics. For the 3D human metastatic lymph node dataset, we used all 16 unique cell types available, providing 256 unique cells pairs (from 16^2^) as input. For the 3D mouse brain cortex dataset, we used the 11 cell types available, providing 121 cell pairs (from 11^2^) as input. For the 3D mouse hypothalamus dataset, we used the 9 cell types available, providing 81 cell pairs (from 9^2^) as input. Finally, we used the corresponding 2D SPIAT metrics (or derived equivalent 2D metrics) on each of the slices that make up each 3D dataset and compared the results from both 3D and 2D analyses using the percentage difference. Percentage difference was calculated as

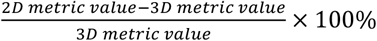

### 5. Comparison of 3D and 2D spatial analysis when comparing between sample groups

Here we aimed to compare the performance of 3D and 2D spatial analysis in identifying differences between sample groups. Specifically, we wanted to identify the false positive and false negative rate of identifying differences between pairs of sample groups in matched 3D and 2D spatial data.

#### 5.1. 3D and 2D simulation generation for dataset comparison between sample groups

We defined a <u>**sample group**</u> as 20 3D tissue simulations (i.e. 20 samples) each generated with spaSim-3D using parameters from the same range (Table S9). A sample group represents ‘biological replicates’ from a group of observations.

A <u>**set**</u> was defined as two sample groups. To test the False Positive Rate (FPR), we generated sets where both sample groups were generated using parameters from the same range, and we would anticipate no significant differences between both groups beyond what is expected by chance (Table S9). To test the False Negative Rate (FNR), we generated groups using different sets of parameter values, where we would anticipate significant differences more often than what is expected by chance (Table S9). Furthermore, in this case, since we wanted to explore the effects of individual parameters on dataset comparisons, only one parameter was changed between the two groups within a set.

Finally, a <u>**collection**</u> was defined as 200 sets. All sets in a collection would be generated using the same parameter ranges (Table S9) used to generate each group within the sets. If the sets were testing FPR, this generated a FPR collection (FPRc) and if the sets were testing FNR, this generated a FNR collection (FNRc). To also test the impact of tissue complexity on the FNR and FPR, simulations with different tissue structure were used when generating the collections. The collections we generated formed Simulated Dataset 3, adding to the simulated datasets generated in section 3.1.

As done previously to obtain matched 2D simulated data for each simulation, we followed the virtual sectioning method described in section 3.2. We randomly selected one 2D slice for each 3D simulation, to mimic what occurs when generating spatial –omics data where a single section is used for profiling.

#### 5.2. Statistical comparisons using Simulated Dataset 3

To compare between each sample groups, we used a two-sided unpaired Wilcoxon test on each SPIAT-3D metric used. This was repeated for the 200 sets within the collection, yielding 200 p-values for each metric, and finally this process was repeated for each collection in Simulated Dataset 3. A similar procedure was followed with the corresponding 2D spatial metrics to the matched 2D sections.

### 6. Performance of 3D and 2D spatial analysis in capturing biological information in time course data

To determine if 2D spatial metrics could accurately capture similar biological trends identified when applying the 3D spatial metrics, we used two sets of publicly available 3D datasets with time course data.

#### 6.1. Publicly available datasets

##### 6.1.1. Mouse embryos 3D dataset

We obtained mouse embryo tissue data from Qiu et al.,^11^ generated using spateo. This included two samples: a mouse embryo sample 9.5 hours and one at 11 hours after birth. We accessed the publicly available H5AD file and extracted the x, y, z coordinates and cell type identity of every cell across the mouse embryo samples. The 9.5 hour embryo sample contained 646,893 cells across 70 slices. The 11 hour embryo sample contained 6,993,667 cells across 84 slices. All slices in the 9.5 hour and 11 hour embryo sample were profiled with spateo.

We identified 6 cell types which overlapped between the 9.5 hour and 11 hour mouse embryo samples (neuroectoderm-derived cells, neural crest cells, primitive erythroid cells, endothelium, cranial mesoderm, placode and neural crest-derived cells). This provided 36 cell pairs (from 6^2^), which were used for subsequent analysis with SPIAT-3D and SPIAT.

##### 6.1.2. *Drosophila* embryo and larvae 3D dataset

We derived a set of 3D tissues from Wang et al.,^12^ who used Stereo-seq to generate spatiotemporal transcriptomic information in a developing *Drosophila*. This included two 3D tissue samples from *Drosophila* during its embryo stage (14–16 hour and 16–18 hour after egg laying), and three 3D tissue samples from *Drosophila* during all three stages of larvae, for a total of 5 consecutive stages of *Drosophila* development.

Each sample was publicly available as separate H5AD files, and each contained the x, y, z coordinates and individual tissue type identity of every cell. The 14-16 hour embryo sample contained 15,295 cells and 16 sections, the 16-18 hour embryo sample contained 14,634 cells and 14 sections, the first larvae stage contained 17,787 cells and 23 sections, the second larvae stage contained 64,658 cells and 21 sections, and finally, the third larvae stage contained 43,310 cells and 16 sections.

We obtained the five distinct individual tissue types (CNS, epidermis, carcass, muscle, midgut, fat body) that overlapped between the five Drosophila stages. This provided 25 cell pairs (from 5^2^) for analysis with SPIAT-3D and SPIAT.

#### 6.2. Comparison of the performance of 3D and 2D spatial analysis when extracting biologically meaningful information

We applied SPIAT-3D metrics with the same parameters used in Table S8 (apart from the cell type inputs) to the mouse embryo and *Drosophila* datasets, as well as the corresponding 2D spatial metrics on the matched 2D data.

Here our goal was to assess whether 2D spatial analysis could capture biological *trends* across time series samples in a manner comparable to 3D metrics, rather than comparing the values of spatial metrics as in previous sections.

To focus on trends across samples, we performed a simple linear transformation of 3D and 2D spatial metrics so they shared the same starting point, which would be used as reference for rest of the samples. For the mouse embryo samples, the spatial metrics from the 9.5 hour embryo were used as the reference: all 3D values were shifted using subtraction such that the 9.5 hour mouse embryo equalled 0, and the same subtraction shift was applied to 11 hour embryo 3D metric values. 2D metrics from the 9.5 hour mouse embryo were transformed using subtraction to a median of 0, and this subtraction shift was applied to the 11 hour mouse embryo 2D values. Likewise, for the *Drosophila* samples, the spatial metrics from the first *Drosophila* stage were used as a baseline: all 3D values were shifted using subtraction such that the 14 - 16 hour *Drosophila* embryo equalled 0, and the same subtraction shift was applied to the 3D metric values of the rest of the samples. Similarly, for the 2D metrics values obtained from the 14 – 16 hour *Drosophila* embryo were also transformed using subtraction to a median of 0, and this subtraction shift was applied to the 2D values of the rest of the samples.

Next, we assessed and quantified whether 2D spatial metrics reproduced the trends observed with the 3D metrics across samples, or whether they systematically overestimated, underestimated, or exhibited inconsistent behaviour even after transformation. To do this, we devised similar but individualised approaches for the mouse embryo and *Drosophila* datasets.

For the mouse embryo samples, we examined how the 11 hour mouse embryo 3D and 2D values compared to the 9.5 hour mouse embryo, which was used as the reference, and developed the following definitions:

1. If the 3D metric value for the 11 hour mouse embryo was contained within the first and third quartiles of the 2D metric values for the 11 hour mouse embryo, we defined the 2D metric values as *‘consistent’* with the ground truth (i.e. 3D metric value).
2. If the 3D metric value for the 11 hour mouse embryo was smaller than the first quartile of the 2D metric slices values for the 11 hour mouse embryo, we defined the 2D metric values as *‘overestimating’* the ground truth (i.e. 3D metric value). Here, the 3D metric value and median 2D metric value for the 11 hour mouse embryo would be following the same trend – they both increased relative the 9.5 hour mouse embryo, or both decreased relative to the 9.5 hour mouse embryo.
3. If the 3D metric value for the 11 hour mouse embryo was greater than the third quartile of the 2D metric values for the 11 hour mouse embryo, we defined the 2D metric values as *‘underestimating’* the ground truth (i.e. 3D metric value). Here, the 3D metric value and median 2D metric value for the 11 hour mouse embryo followed the same trend – they both increased relative the 9.5 hour mouse embryo, or both decreased relative to the 9.5 hour mouse embryo.
4. In all other cases, the 2D metric values were classified as *‘inconsistent’*. In this case, the 3D metric value was greater while the median 2D metric value was lower relative to the 9.5 hour mouse embryo, or vice versa.

For the *Drosophila* samples, we used a different approach as there were 5 samples to compare. Since the 3D metric values and median 2D metric value for the 14 – 16 hour *Drosophila* embryo serve as the reference points, we focused on the trends of the remaining 4 samples of the series. We developed the following definitions:

1. The 2D metric trends were *‘consistent’* with the ground truth (i.e. 3D metric) trends, if the 3D metric value was within the first and third quartile of the 2D metric slices values in at least 3 of the 4 remaining samples.
2. The 2D metric trends were *‘overestimating’* the ground truth (i.e. the 3D metric) trends if the 3D metric value was smaller than the first quartile formed by the 2D metric slices values in at least 3 of the 4 remaining samples.
3. The 2D metric trends were *‘underestimating’* the ground truth (i.e. 3D metric) trends if the 3D metric value was greater than the third quartile of the 2D metric slices values in at least 3 of the 4 remaining samples.
4. The 2D metric value trends were *‘inconsistent’* compared to the ground truth (i.e. 3D metric) trends if it did not fall under any of these previous categories. In this scenario, the 3D metric value varied between being between the first and third quartiles, below the first quartile or above the third quartile across the 4 remaining samples.

For a given metric, we determined which category the 2D metric trend fell under (consistent, overestimating, underestimating or inconsistent) for each input cell pair (36 cell pairs for the mouse embryo dataset and 25 cell pairs for the *Drosophila* dataset). We then aggregated the results by computing the proportion of cell pairs falling into each category for that metric, and repeated this for all metrics.

## Code availability

The code for the spaSim-3D tool is available at https://github.com/TrigosTeam/spaSim-3D. The code for the SPIAT-3D tool is available at https://github.com/TrigosTeam/SPIAT-3D. The code to reproduce the work presented is available at https://github.com/TrigosTeam/spaSim-3D_SPIAT-3D_analysing_individual_datasets, https://github.com/TrigosTeam/spaSim-3D_SPIAT-3D_comparing_datasets, https://github.com/TrigosTeam/spaSim-3D_SPIAT-3D_paper_figures.

## Data availability

All biological data used in this work was obtained from publicly available sources. The data for the 3D human colorectal cancer dataset was obtained from https://zenodo.org/records/7554924; the data for the 3D human metastatic lymph node dataset was obtained from GSE251926; the data for the 3D mouse embryo dataset was obtained from https://spateodata.aristoteleo.com; the data for the 3D mouse brain dataset was obtained from https://www.dropbox.com/scl/fo/7fxdv5rmnslhaffsczqbg/AGV_LSSObDifq2FfhtbMwqk?rlkey=qoyw13ddac65n3ui2hzaw9ber&e=1&dl=0%2F and the 3D drosophila larvae and embryo dataset was obtained from https://db.cngb.org/stomics/flysta3d/download/.

## Acknowledgements

We would like to thank Prof. Terry Speed and A/Prof. Ute Hahn, the Centre for Advanced Histology & Microscopy (CAHM) (RRID:SCR_025432) and the Peter Mac Research Computing Facility. This work was supported by an NHMRC Ideas grant 2020149, a L’Oreal-UNESCO Fellowship and a Prostate Cancer Foundation TACTICAL Award, all awarded to AST. AST is a Prostate Cancer Foundation Young Investigator, class of 2021. This work was supported by resources provided by the Pawsey Supercomputing Research Centre’s Setonix Supercomputer (https://doi.org/10.48569/18sb-8s43), with funding from the Australian Government and the Government of Western Australia. We also thank Peter Sorger and his team for providing updated versions of the publicly available human colorectal cancer dataset used in this study.

## Author contributions

DL designed and performed all simulations and analyses, wrote the code, designed tools, interpreted results, wrote and edited the manuscript. DIK designed analysis, interpreted results and supervised the work. AST conceived and supervised the work, developed research questions, designed analyses, interpreted results, wrote and edited the manuscript.

