## Supplementary Figures for "2D spatial tissue analysis often misrepresents true biological patterns of 3D tissues"

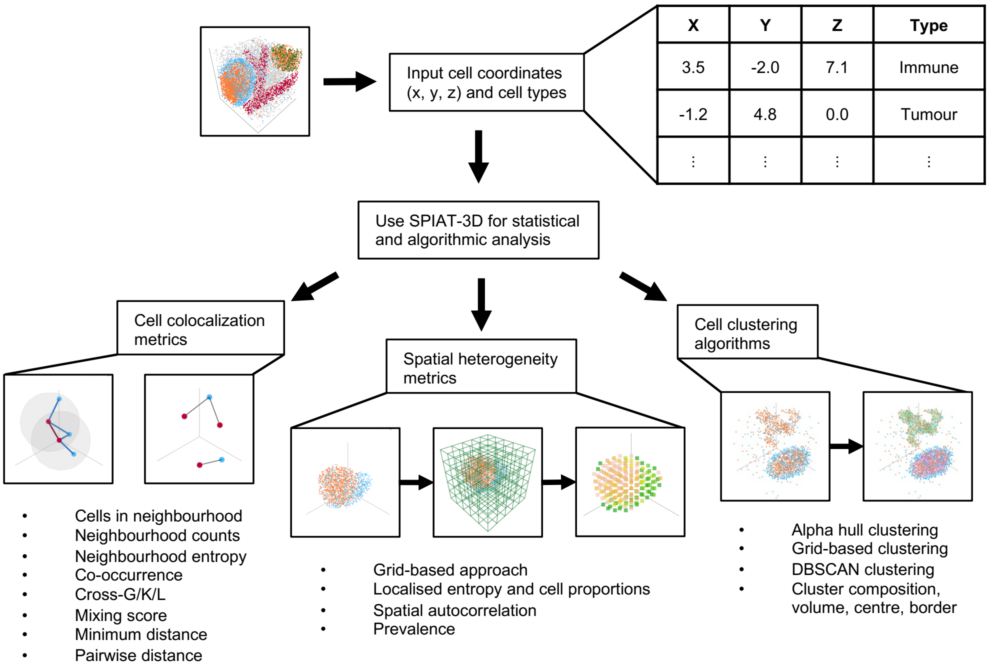


**Supplementary Figure S1. Overview of SPIAT-3D.** SPIAT-3D calculates a range of spatial analysis metrics, including cell colocalization metrics, spatial heterogeneity metrics and cell-clustering algorithms. SPIAT-3D is based on SPIAT, which enabled similar calculations of 2D spatial data. SPIAT-3D also introduces new metrics such as neighbourhood counts and neighbourhood entropy, and adapts existing 2D spatial metrics from other packages, such as co-occurrence from squidpy.[6] As with SPIAT, the starting input of SPIAT-3D is a table with the x, y and z coordinates of cells and categorical cell types.

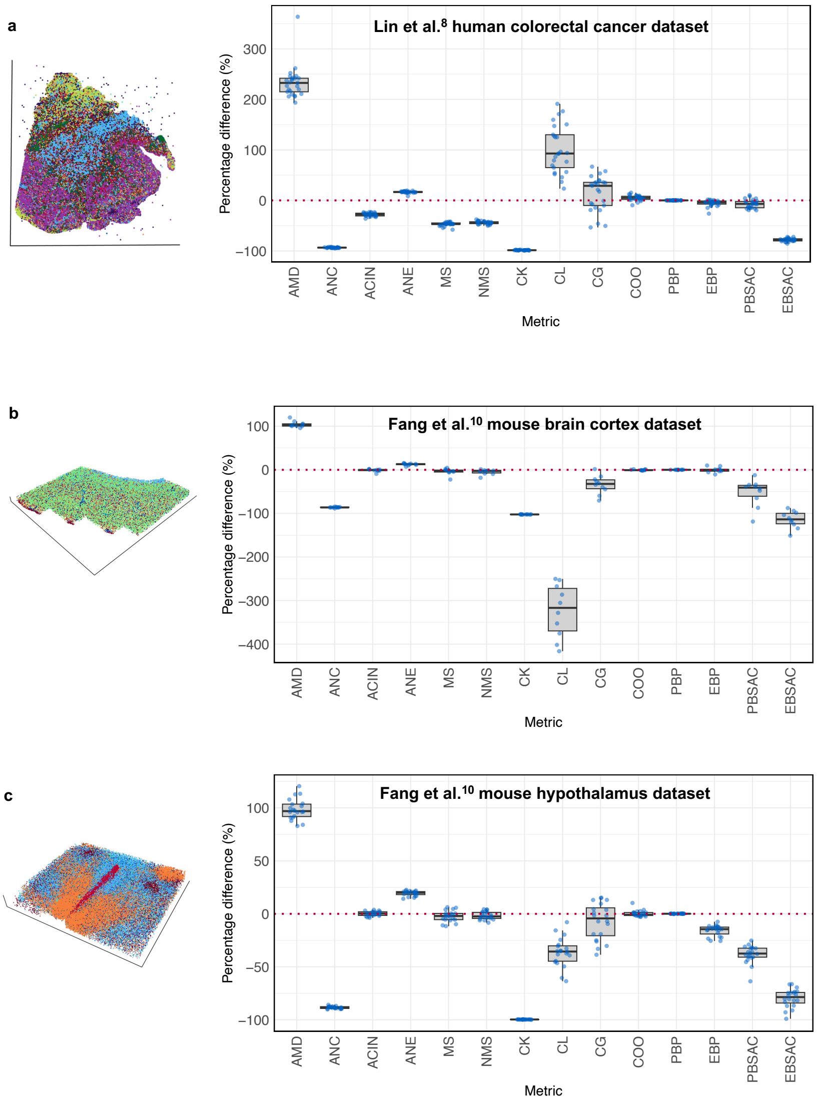


**Supplementary Figure S2. Percentage differences between 3D and 2D metrics in publicly available 3D datasets.** Box plots showing the distribution of median percentage differences between each 3D metric and its equivalent 2D metric. The percentage difference was calculated as: $percentage\ difference = \left(\frac{2D\ value - 3D\ value}{3D\ value}\right) \times 100\%$. 3D metrics were applied to each 3D dataset while 2D equivalent metrics were applied to each extracted slice from the same 3D dataset. Multiple cell pairs (e.g. Tumour/Tumour, Tumour/Immune, etc.) were used as inputs for the 3D and 2D metrics. For each slice, the percentage difference between the 3D and 2D metric values was computed for every cell pair. The median percentage difference across all cell pairs was then taken for that slice. These medians for each slice form the points shown in the box plots. Results shown for the **(a)** human colorectal cancer sample generated with CyCIF by Lin et al.,[8] **(b)** mouse brain cortex sample generated with MERFISH by Fang et al.[10] and **(c)** mouse hypothalamus sample generated with MERFISH by Fang et al.[10] Including the human metastatic lymph node sample generated with Open-ST by Schott et al.[9] shown in Figure 1b, the AMD, CL and PBSAC metrics were the top three metrics with highest discrepancy across all four datasets (the range of the percentage difference was 32.45-363.61%, -416.21-191.15%

and -118.71-15.96%, respectively), while CK and PBP showed the lowest discrepancy (-103.03- -97.70%, -0.29-0.35% respectively).

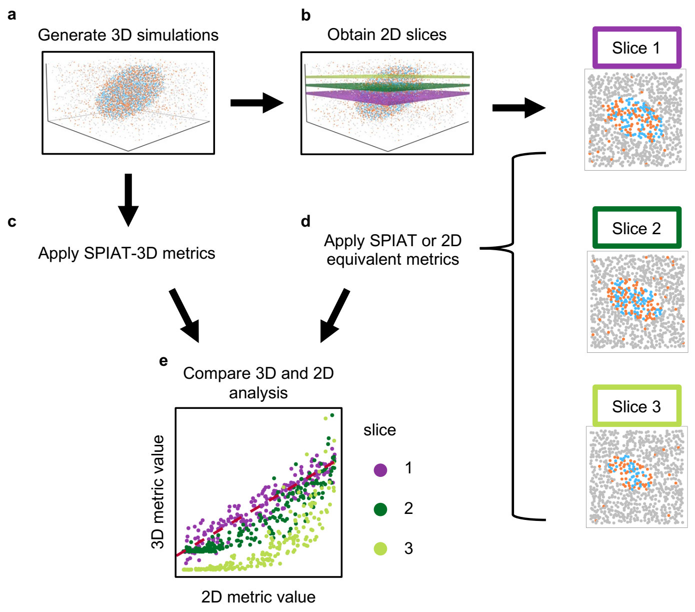


**Supplementary Figure S3. Virtual tissue sectioning and comparison between 3D and 2D spatial metrics results.** **(a)** The starting point is a 3D sample from a biological dataset or simulated with spaSim-3D. **(b)** 2D slices are extracted from the x-y plane from each 3D simulation. This process aims to recapitulate what occurs in a laboratory setting when tissues are sectioned for the generation of spatial –omics data. Each slice had a width of 10 units, corresponding to the 'minimum distance between cells' parameter used when generating the background cells of each simulation, ensuring that each slice contained, roughly, a single layer of cells. **(c)** SPIAT-3D metrics are applied to each 3D sample. **(d)** 2D equivalent metrics are applied to each 2D slice. **(e)** Results from 3D and 2D analyses are compared between matched 3D samples and 2D slices.

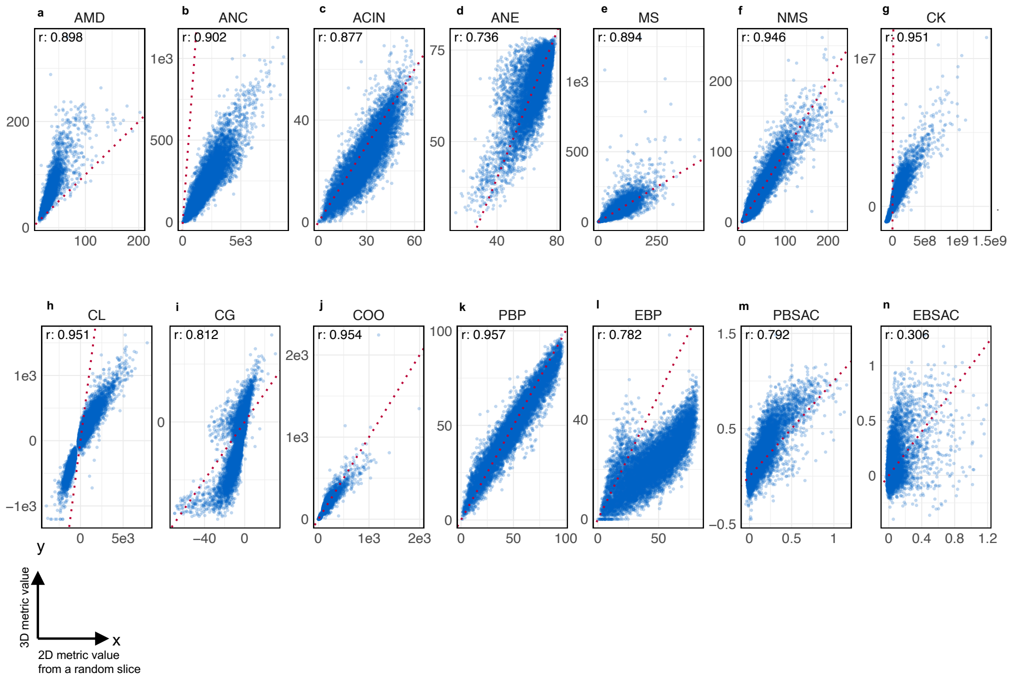


**Supplementary Figure S4. Comparison of 3D and 2D metrics applied to diverse 3D simulations and one random 2D slice from each simulation.** Scatter plots illustrating the relationship between each 3D metric and its equivalent 2D metric. 10,000 3D simulations were generated by varying both categorical and continuous parameters such as cluster arrangement, shape and background noise. 13 2D slices were taken from each 3D simulation, with one slice randomly selected. Each scatter plot is based on 10,000 data points, representing one 2D slice from each of 10,000 3D simulations. Spearman correlations were preformed to assess the strength of the 2D-3D relationship. **(a)** AMD. **(b)** ANC. **(c)** ACIN **(d)** ANE **(e)** MS **(f)** NMS **(g)** CK **(h)** CL **(i)** CG **(j)** COO **(k)** PBP **(l)** EBP **(m)** PBSAC **(n)** EBSAC.

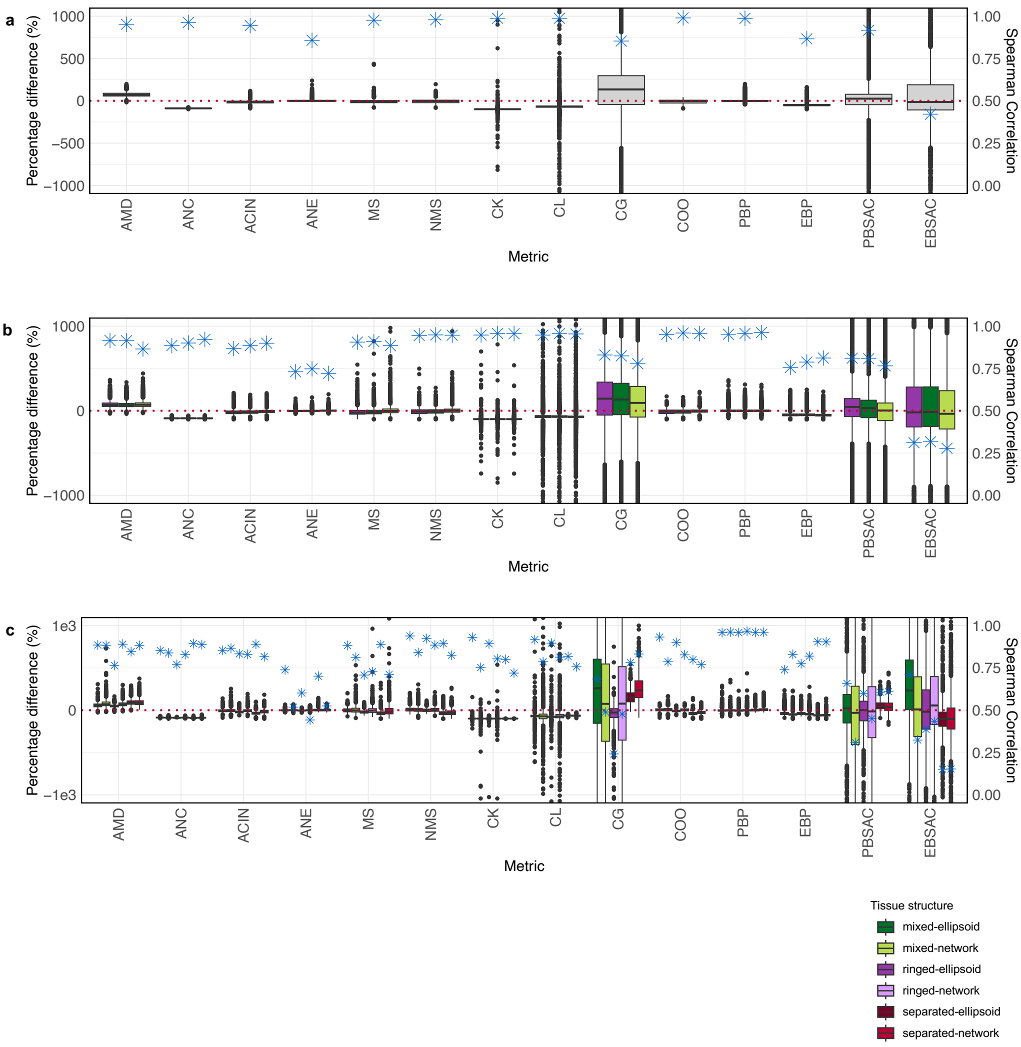


**Supplementary Figure S5. Comparing 3D and 2D metrics applied to diverse 3D simulations and selections of 2D slices from each simulation.** Box plots showing the distribution of percentage differences between each 3D metric and its equivalent 2D metric with spearman correlations (shown as a blue star) between 3D and 2D metric values. The percentage difference was calculated as: $percentage\ difference = \left(\frac{2D\ value - 3D\ value}{3D\ value}\right) \times 100\%$. 10,000 3D simulations were generated by varying both categorical and continuous parameters such as cluster arrangement, shape and background noise. 3D metrics were applied to each simulation. **(a)** 13 2D slices were taken from each 3D simulation. 2D metrics were applied to each slice and an average was taken for each metric and simulation. Percentage difference and spearman correlation were calculated between the 3D metric values and the 2D metric averages. Data points with percentage difference below -1000% or above 1000% are not shown for visualisation and can be found in table S10. **(b)** 3 2D slices were taken from each simulation and 2D metrics were applied to each slice (purple: middle slice, dark green: upper slice, light green: uppermost slice). Percentage difference and spearman correlation were calculated between the 3D metric values and the 2D metric values for each slice. Data points with percentage difference below -1000% or above 1000% are not shown for visualisation and can be found in table S11. **(c)** Simulations were subsetted based on their tissue structure (mixed-ellipsoid, mixed-network, ringed-ellipsoid, ringed-network, separated-ellipsoid, separated-network). 13 2D slices were taken from each 3D simulation and 2D metrics were applied to one randomly selected slice. Percentage difference and spearman correlation were calculated between the 3D metric values and the 2D metric values for each tissue structure. Data points with percentage difference below -1000% or above 1000% are not shown for visualisation and can be found in table S12.

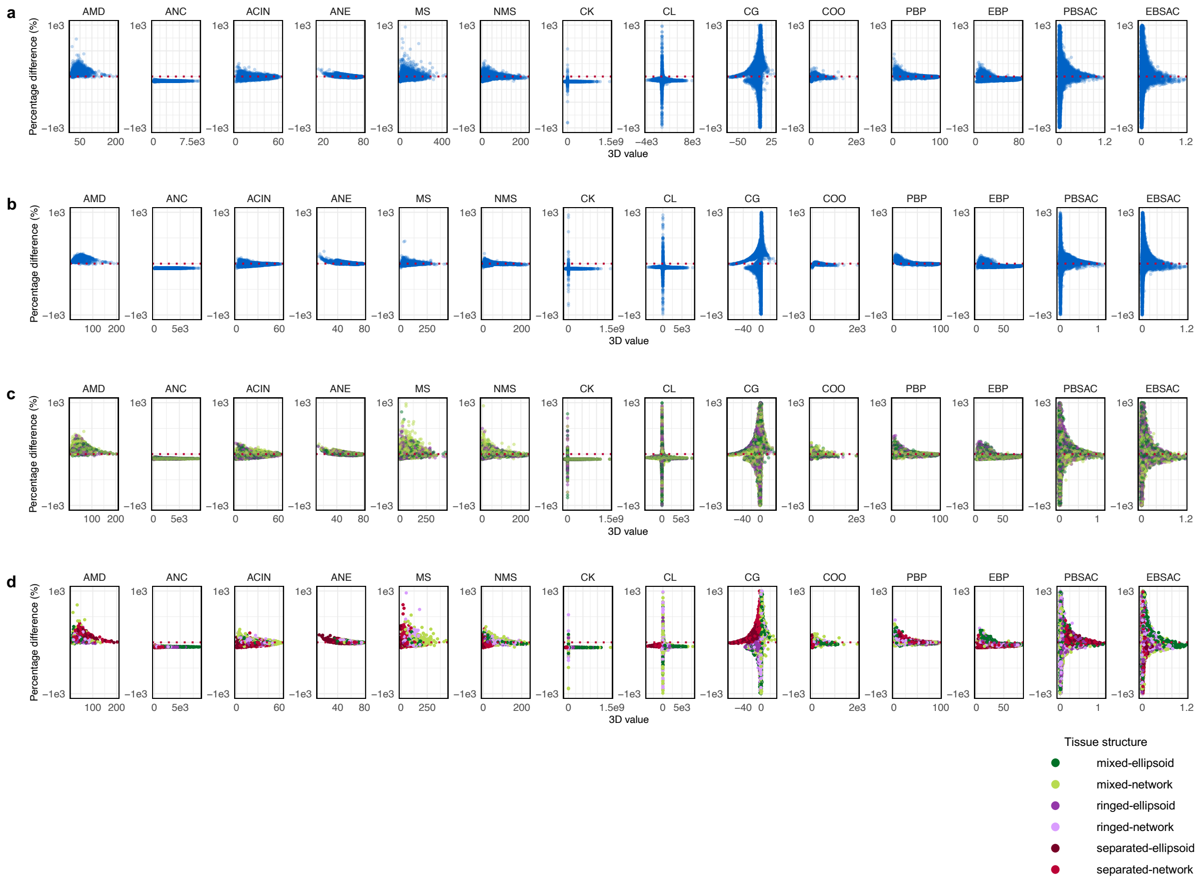


**Supplementary Figure 6. Percentage differences between 3D metrics and 2D metrics for all 14 metrics.** Data shown is the same as Figure 1d and Supp. Fig. S5, but plotting against the 3D values. **(a)** Random slice. Data points with percentage difference below -1000% or above 1000% are not shown for visualisation and can be found in table S12. **(b)** Average across 13 slices. Data points with percentage difference below -1000% or above 1000% are not shown for visualisation and can be found in table S10. **(c)** Values for the middle slice (purple), upper slice (dark green), uppermost slice (light green). Data points with percentage difference below -1000% or above 1000% are not shown for visualisation and can be found in table S11. **(d)** Values coloured based on their tissue structure (mixed-ellipsoid, mixed-network, ringed-ellipsoid, ringed-network, separated-ellipsoid, separated-network). Data points with percentage difference below -1000% or above 1000% are not shown for visualisation and can be found in table S12.

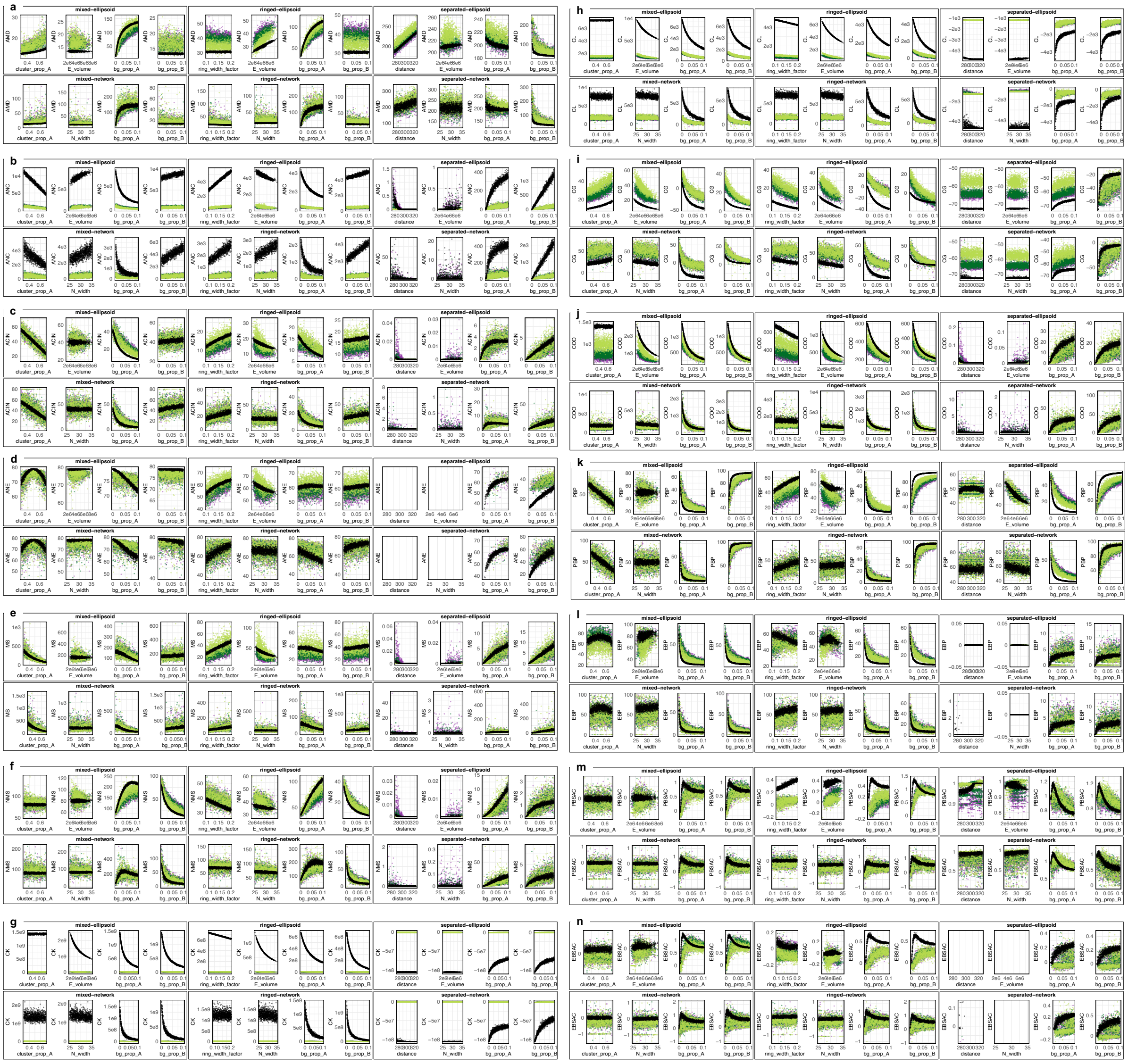

**Supplementary Figure S7. Simulations of varying parameters showing associations between 3D and 2D metrics and the parameters.** Scatter plots showing the relationship between each metric and various continuous variable parameters for the 6 categorical parameter sets: mixed-ellipsoids, mixed-networks, ringed-ellipsoids, ringed-networks, separated-ellipsoids, separated-networks. 1000 3D simulations were generated by systematically varying a single continuous variable parameter (cluster_prop_A, ring_width_factor, distance, bg_prop_A, bg_prop_B, E_volume, or N_width) while keeping others constant. Description for each continuous variable parameter can be found in Table S6. 3D spatial metrics were applied to each simulation. 3 2D slices were taken from each simulation and the corresponding 2D metric was applied to each slice. Each scatter plot is based on 4000 data points, representing the 2D metric value obtained from the 3 2D slices from each of the 1000 3D simulations (purple: middle slice, dark green: upper slice, light green: uppermost slice), along with 1000 additional points representing the 3D metric value obtained from the original 3D simulation (shown by black points). **(a)** AMD. **(b)** ANC. **(c)** ACIN **(d)** ANE **(e)** MS **(f)** NMS **(g)** CK **(h)** CL **(i)** CG **(j)** COO **(k)** PBP **(l)** EBP **(m)** PBSAC **(n)** EBSAC. The distance between structures (distance), the volume of ellipsoid structures (E_volume) and the width of network edges (N_width) resulted in uncorrelated 3D and 2D values in ANC, ACIN, MS, NMS, COO, PBP and PBSAC. Disruption of the 3D and 2D association also occurred in simulations with mixed-ellipsoid structures for EBP, separated tissue structures and with varying background proportion of cells (bg_prop_A and bg_prop_B) for EPB and EBSAC, in cases of ringed-ellipsoid structures for PBSAC, in cases of extensive cell mixing (cluster_prop_A) when using the ANE.
