## Supplementary Notes for "2D spatial tissue analysis often misrepresents true biological patterns of 3D tissues"

#### Supplementary Note 1
#### SPIAT-3D

##### 1.1. Characterising cell colocalization

One category of spatial metrics to help quantify the spatial relationship between different cells within a tissue are cell colocalization metrics. Here we outline the various strategies used in SPIAT-3D to measure cell colocalization.

###### 1.1.1. Distance-based metrics

Distance-based metrics can be used to examine how cells are interacting by quantifying how far they are from each other. Cell types at a closer distance to each other indicate potential communication between these cells. Two methods for calculating cell-cell distances include pairwise distance and minimum distance.

###### 1.1.1.1. Pairwise distance between cells

For a given pair of cell types specified by the user, for example tumour cells and immune cells, pairwise distance analysis involves computing and tabulating the distance between each tumour cell and each immune cell (Figure N1.1a).[1] To do this, we used the '*Negative Distance Matrix*' function from the apcluster package[13] and inputted the x, y, z coordinates of all cell types of interest. This outputs a distance matrix, showing the distance for each cell-cell pair, which is further processed for analysis and plotting.

###### 1.1.1.2. Minimum distance between cells

For a given reference cell type and target cell type specified by the user, minimum distance analysis involves starting at each reference cell, calculating the distance to the closest target cell and tabulating these results (Figure N1.1b).[1] To implement this, we used the '*Nearest Neighbour Search*' function from the RANN package.[14] Inputting the x, y, z coordinates of all reference cells and target cells into this function yields all closest reference-target pairs and their corresponding minimum distances.

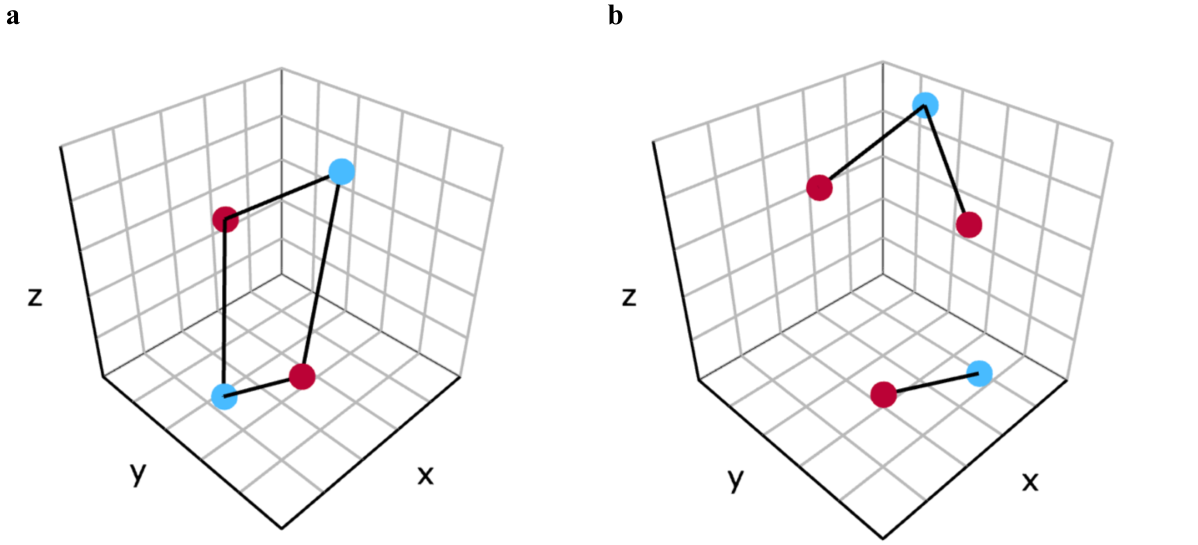


**Figure N1.1. Visualisation of distance-based metrics. (a)** Pairwise distances between cells, showing connections between red and blue cells. **(b)** Minimum distances between reference cells (red) and target cells (blue).

##### 1.1.2. Neighbourhood counts

The neighbourhood counts (NC) metric can be used to quantify how densely populated certain cell types are around another cell type. It can provide insights into whether a particular cell type tends to cluster around another cell type. In SPIAT, many cell colocalization are first calculated by first choosing a reference cell type and a set of target cell types and determining the 'number of interactions between the reference cell and target cells' at a chosen distance. Geometrically, this is the same as counting the number of target cells found in the circle drawn around each reference cell, where the radius of this circle equals the chosen distance. For SPIAT-3D, this concept has been adapted. Instead of a circle, a sphere is drawn around each reference cell and the number of target cells found in each sphere is quantified (Figure N1.2). If multiple target cell types are selected, a separate NC value corresponding to each target cell type would be calculated for each reference cell. To implement the NC metric in SPIAT-3D, we used the '*Fixed Radius Nearest Neighbors*' function from the dbscan package[15] which takes the x, y, z coordinates of the reference cells and target cells, as well as the sphere radius or chosen distance as input, and outputs the NC value for each reference cell.

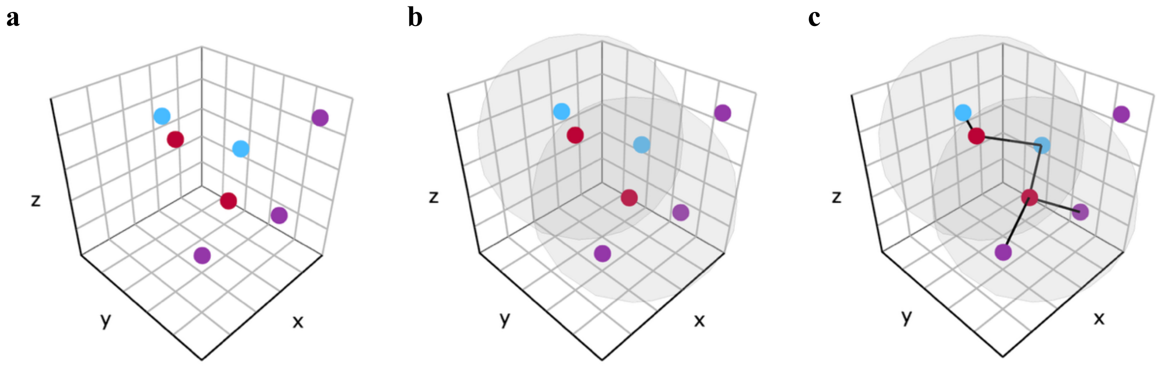


**Figure N1.2.** Visualisation of NC. **(a)** A random set of cells in a 3D plane. Reference cells are red, and target cells are blue and purple. **(b)** Spheres with radius equal to the chosen distance are drawn around each reference cell. **(c)** Target cells inside each sphere are defined as interacting with the reference cell, shown by the blue and purple edges. The number of interactions with each reference cell are used to calculate NC values for each target cell type.

The equation for NC is as follows:

(1) $$\boldsymbol{Neighbourhood\ counts\ (NC) = n_{reference\ to\ target\ interactions}}$$

Where:

- $n_{reference\ to\ target\ interactions}$ is the number of interactions between a particular reference cell and target cells, which is equivalent to the number of target cells found in the sphere drawn around a particular reference cell.

##### 1.1.3. <u>Cells in neighbourhood</u>

NC quantifies the absolute number of target cells interacting with a reference cell.[1] Cells in neighbourhood (CIN) is a metric used in SPIAT that extends the NC metric by measuring the proportion of those interactions: specifically, the fraction of interacting cells that are target cells, relative to the number of target and reference cells interacting with the reference cell. By doing so, we can analyse the relative abundance of a chosen target cell type with respect to a reference cell type and potentially find patterns that might not be apparent when considering absolute counts alone.

The equation for CIN is as follows:

(2) $$\boldsymbol{Cells\ in\ neighbourhood\ (CIN) = \frac{NC_{target}}{NC_{target}+NC_{reference}}}$$

Where:

- $NC_{target}$ is the NC value of a chosen target cell type for a particular reference cell.
- $NC_{reference}$ is the NC value of the chosen reference cell type for a particular reference cell.

##### 1.1.4. <u>Neighbourhood entropy</u>

For a tissue containing different cell types, entropy measures the level of disorder in the proportion of cell types in the tissue. Low disorder and hence, low entropy, occurs when a tissue contains a high proportion of a particular cell type. On the other hand, when there is a roughly equal mix of different cell types in a tissue, disorder and entropy are high. Applying entropy to the entire tissue, we can quantify the heterogeneity of cell types, providing a measure of how evenly or unevenly they are distributed across the tissue. This concept of entropy can be further expressed using CIN, which we termed 'neighbourhood entropy (NE)'. For a reference cell type, target cell type and radius selected by the user, if the CIN value of the reference and target cell type is similar, NE is high. However, if the CIN value of the reference and target cell type differs greatly, when either the reference or target cell dominates in proportion, NE is low (Figure N1.3). Through this, we can determine whether a particular cell type is generally surrounded by one cell type or an equal mixture of cell types.

**a** **b**

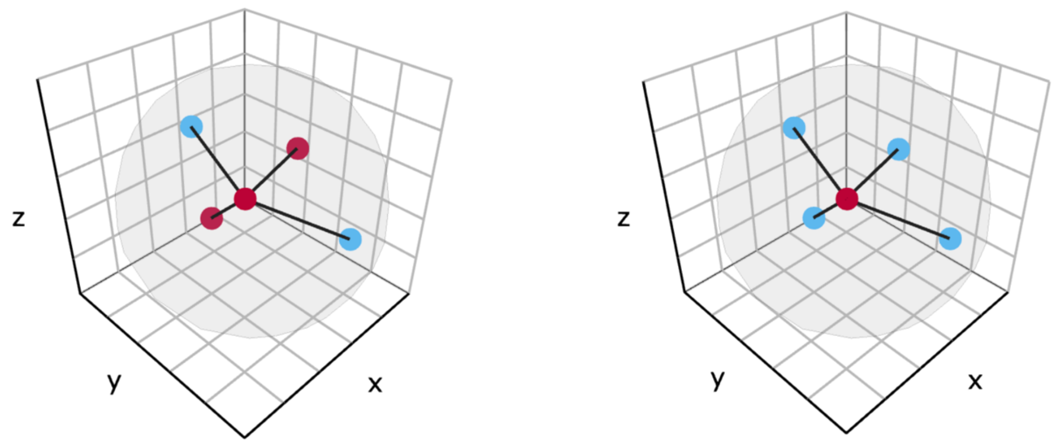


**Figure N1.3. Visualisation of NE. (a)** High NE as there is balance in the proportion of reference and target cells (red and blue) interacting with the reference cell (red). **(b)** Low NE as the proportion of reference and target cells interacting with the reference cells is dominated by the target cells.

We adapted the general entropy equation to derive the equation for NE:

**(3)** $$\boldsymbol{Entropy} = -\sum_{i=1}^{n} \boldsymbol{p(X_i)} \log_2 \boldsymbol{p(X_i)}$$

Where:

- $X_i$ is the ith event of a system.
- $p(X_i)$ is the proportion of $X_i$ in the whole system.

Replacing $p(X_i)$ with CIN values for reference and target cells yields the neighbourhood entropy formula:

**(4)** $$\boldsymbol{CIN_{reference} = 1 - CIN_{target}}$$

**(5)** $$\boldsymbol{Neighbourhood\ entropy\ (NE) = -(CIN_{target} \log_2(CIN_{target}) + CIN_{reference} \log_2(CIN_{reference}))}$$

Where:

- $CIN_{target}$ is the *CIN* value for the target cell type.
- $CIN_{reference}$ is the *CIN* value for the reference cell type.

##### 1.1.5. Mixing scores and normalised mixing scores

The mixing score (MS) metric quantifies the degree of colocalization between a reference cell type and a target cell type.[2] A high MS indicates frequent interactions between the two, while a low MS suggests limited interaction and a tendency for the reference cell type to cluster with itself. However, variations in the number of reference and target cells present in different tissues can make MS difficult to interpret. Adapting MS, Normalised mixing score (NMS) considers the number of reference and target cells present and can be benchmarked against a CSR distribution.[1] An NMS value of 1 suggests frequent interactions between the reference and target cell types that follow a CSR distribution (Figure N1.4). Similar to the NC, CIN and NE metric, both MS and NMS are calculated by finding the number of target cells inside the sphere drawn around the reference cell. However, when using NC, CIN and NE, a separate value is calculated for each reference cell, but for MS and NMS, the number of target and reference cells found inside each reference cell's sphere is instead summed or totalled across the entire tissue. This results in a single MS or NMS value for the chosen distance.

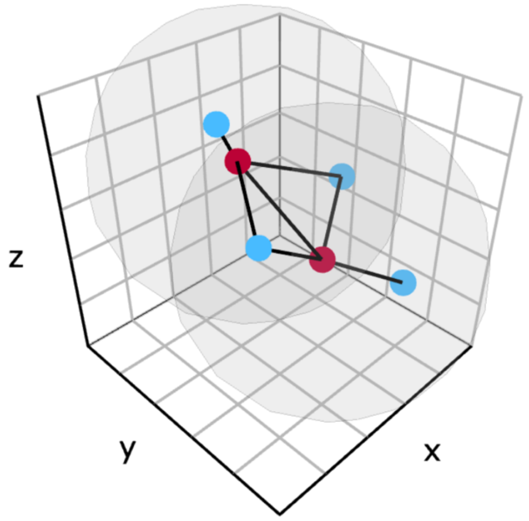


**Figure N1.4. Visualisation of MS and NMS.** The number of reference cells (red) and target cells (blue) interacting with or in the sphere of each reference cell is totalled and used to calculate MS and NMS.

Equations for MS and NMS are as follows:

(6) $$\boldsymbol{Mixing\ score\ (MS)} = \frac{\sum \boldsymbol{NC_{target}}}{\sum \boldsymbol{NC_{reference}}}$$

(7) $$\boldsymbol{Normalised\ mixing\ score\ (NMS)} = \frac{\boldsymbol{n_{reference}}}{\boldsymbol{2 \times n_{target}}} \times \boldsymbol{MS}$$

Where:

- $\sum \boldsymbol{NC_{target}}$ is the sum of NC values of a chosen target cell type across all reference cells.
- $\sum \boldsymbol{NC_{reference}}$ is the sum of NC values of the chosen reference cell type across all reference cells.
- $n_{reference}$ is the number of reference cells in the tissue.
- $n_{target}$ is the number of target cells in the tissue.

##### 1.1.6. Cross K-function point estimate

The cross K-function (CK) quantifies the levels of clustering or repulsion between different types of points.[3] Applying it to cells in a tissue, similar to NC, the CK can provide information of whether a particular cell type tends to cluster around another cell type or repel it. Unlike NC and other metrics, the CK considers the tissue area or volume, allowing for a direct comparison between the observed clustering or repulsion of two cell types and the expected clustering or repulsion if they were randomly arranged following a CSR distribution.

For a given reference cell type, target cell type and radius specified by the user, an observed CK greater than the expected CK indicates clustering between the reference and target cell, while the opposite indicates repulsion (Figure N1.5). One thing to note is that by using a single radius, we calculate a point estimate of the CK. However, the CK is generally calculated over a range of radii, which provides insight into clustering or dispersion patterns at different distances. This concept of using a range of radii is covered in section 1.1.10.

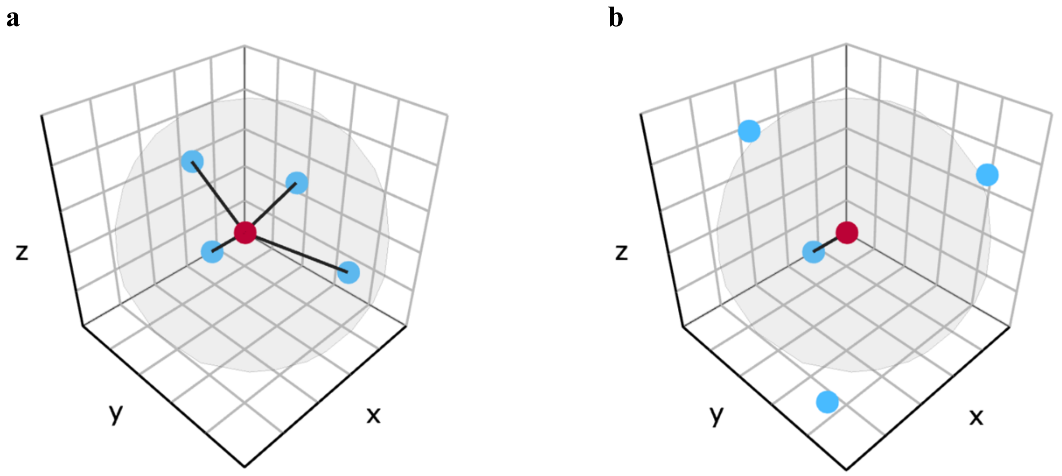


**Figure N1.5. Visualisation of CK point estimate. (a)** High observed CK point estimate as all target cells (blue) are interacting with the reference cell (red) **(b)** Low observed CK point estimate as few target cells are interacting with the reference cell.

Equations for the CK are as follows:

(8) $$\boldsymbol{Observed\ cross\ K - function\ (CK) = \frac{V_{tissue} \times \sum NC_{target}}{n_{reference} \times n_{target}}}$$

(9) $$\boldsymbol{Expected\ cross\ K - function\ (CK) = \frac{4}{3}\pi r^3}$$

Where:

- $r$ is the radius of the sphere drawn around each reference cell. Cells within the sphere are counted as interacting with the reference cell.
- $\sum \boldsymbol{NC_{target}}$ is the sum of NC values of a chosen target cell type across all reference cells.
- $n_{reference}$ is the number of reference cells in the tissue.
- $n_{target}$ is the number of target cells in the tissue.
- $V_{tissue}$ is the volume of the tissue.

1.1.7. Cross L-function point estimate

The cross L-function (CL) is a linearised version of the CK.[4] While the CK follows cubic growth (as shown in equation (**9**)), the CL is transformed so it follows linear growth such that the expected CL is equal to the chosen radius (see equation (**11**)). The CK and CL will show the same trends, but the CL may be preferred as it grows linearly. It is also straightforward to interpret as when compared to a CSR distribution, an observed CL greater than the chosen radius suggests clustering, whilst the opposite suggests repulsion. As is the case for the CK, while a point estimate of the CL at a single radius is possible, the CL is generally calculated over a range of radii, which is detailed in section 1.1.10.

Equations for the CL are as follows:

$$(10)\quad \boldsymbol{Observed\ cross\ L - function\ (CL) = \sqrt[3]{\frac{3}{4\pi} \times observed\ CK}}$$

$$(11)\quad \boldsymbol{Expected\ cross\ L - function\ (CL) = \sqrt[3]{\frac{3}{4\pi} \times expected\ CK} = r}$$

Where:

- $r$ is the radius of the sphere drawn around each reference cell. Cells within the sphere are counted as interacting with the reference cell.

1.1.8. Cross G-function point estimate

The cross G-function (CG) is another cell colocalization metric.[5] Unlike the CK or CL, which consider all target cells within the sphere drawn around each reference cell, the CG focuses only on the nearest target cell, potentially making it more sensitive to small-scale cell-cell interactions. Specifically, the CG gives the proportion of reference cells whose nearest target cell neighbour is within a chosen distance. When compared to a CSR distribution, observed CG values above the expected CG suggests clustering between reference and target cells, while the opposite suggests repulsion. Again, similar to the CK and CL, the CG can be calculated at a single distance threshold, providing a point estimate. However, the CG is more commonly calculated over a range of radii, which is detailed in section 1.1.10.

Equations for the CG are as follows:

$$(12)\quad \boldsymbol{Observed\ cross\ G - function\ (CG) = P(distance\ from\ a\ reference\ cell\ type\ to\ nearest\ target\ cell\ type \leq r)}$$

$$(13)\quad \boldsymbol{Expected\ cross\ G - function\ (CG) = 1 - e^{\left(-\frac{n_{target}}{V_{tissue}} \times \frac{4}{3}\pi r^3\right)}}$$

Where:

- $r$ is the radius of the sphere drawn around each reference cell. Cells within the sphere are counted as interacting with the reference cell.
- $n_{target}$ is the number of target cells in the tissue.
- $V_{tissue}$ is the volume of the tissue.

1.1.9. Co-occurrence

Co-occurrence (COO) is a spatial metric implemented in the squidpy package.[6] It has been adapted here, using the nomenclature seen in previous SPIAT-3D metrics. For a given sphere radius, a COO metric (see equation (14)) is calculated by determining the proportion of target cells found within the sphere of each reference cell, and comparing this to the overall proportion of target cells found within the tissue. If the cells in the tissue followed a CSR distribution, COO would equal 1, as the proportion of target cells around reach reference cell would match the proportion of target cells in the tissue sample. Hence, interpreting co-occurrence is straightforward, as observed COO values greater than 1 indicate clustering between reference and target cells, while values less than 1 indicate repulsion.

The equation for COO is as follow:

$$(14) \quad \boldsymbol{Co-occurrence\ (COO)} = \frac{\sum NC_{target}}{\sum NC_{all}} / \frac{n_{target}}{n_{cells}}$$

Where:

- $\sum \boldsymbol{NC_{target}}$ is the sum of NC values of a chosen target cell type across all reference cells.
- $\sum \boldsymbol{NC_{all}}$ is the sum of NC values of all cell types across all reference cells.
- $n_{target}$ is the number of target cells in the tissue.
- $n_{cells}$ is the number of cells in the tissue.

1.1.10. Gradient-based metrics

NC, CIN, NE, MS, NMS, CK, CL, CG, and COO are all based on the concept of a sphere being drawn around each reference cell, and the number of interactions between reference and target cell, or in other words, the number of target cells found in these spheres is counted. These metrics can be calculated for a single distance/sphere radius – this gives users the ability to choose the type of cellular interactions they are interested in. For example, setting this distance to be slightly larger than the diameter of a cell may implicate cell-cell interactions, while using a much larger distance may provide information about indirect communication between cells.

However, to avoid making an (often arbitrary) decision on the radius, an alternative is to observe how these metrics behave across a gradient of increasing sphere radii (Figure N1.6a), and plotting the output of the chosen metric against this gradient of sphere radii. This is traditionally how the CK, CL and CG are calculated.[3–5]

For MS, NMS, CK, CL, CG, and COO, a single value is outputted for each sphere radius, and the gradient-based graph can be plotted. In contrast, for NC, CIN, and NE, each reference cell has its own corresponding value. Therefore, to plot the gradient-based graph for these metrics, an average value across all reference cells is calculated for each sphere radius.

By calculating these cell colocalization metrics across a gradient, we can understand how the levels of interaction between cells varies at different distances. An example simulation and corresponding gradient-based plot is shown in Figure N1.6.

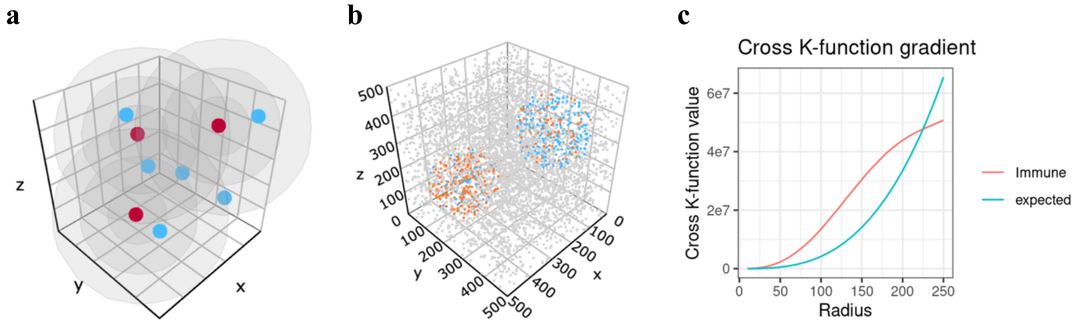


**Figure N1.6. Visualisation and plotting of gradient-based cell colocalization metrics. (a)** Spheres of increasing radii are drawn around each reference cell (red). The number of interactions between reference cells and target cells (blue) at each radius can be used to calculate a chosen metric for each radius. **(b)** Example tissue simulation generated by spaSim-3D containing tumour cells (orange) and immune cells (blue). **(c)** CK gradient plot for the tissue simulation in Figure N1.6b. Observed (tumour cell as reference cell and immune cell as target cell) and expected CK values were calculated for sphere radii increasing from 50 to 250 in increments of 25. Clustering between tumour and immune cells is shown for radius values below ~225, and dispersion is shown for radius values above ~225.

#### 1.2. Characterising spatial heterogeneity

While cell colocalization examines the association between different cells, spatial heterogeneity is defined as the variation in the spatial distribution of cells across a tissue. For example, it can help visualise and quantify the uneven arrangements of tumour cells in a cancer sample.

In SPIAT, spatial heterogeneity in a 2D tissue section is quantified by dividing the tissue into a 2D grid of rectangles, analysing the cellular composition of each rectangle, and applying spatial heterogeneity metrics to quantify the distribution of grid rectangles (Figure N1.7, top row).[1] We have extended this approach to 3D tissue analysis for SPIAT-3D. This adapted method involves breaking down the 3D tissue into a grid of many rectangular prisms, the number of which can be controlled by the user. The cellular composition and spatial location of each rectangular prism are then used as input for spatial heterogeneity metrics in SPIAT-3D (Figure N1.7, bottom row). These metrics have been adapted from SPIAT to consider the x, y, z coordinates of each rectangular prism and innovated to showcase spatial heterogeneity data in a more informative way.

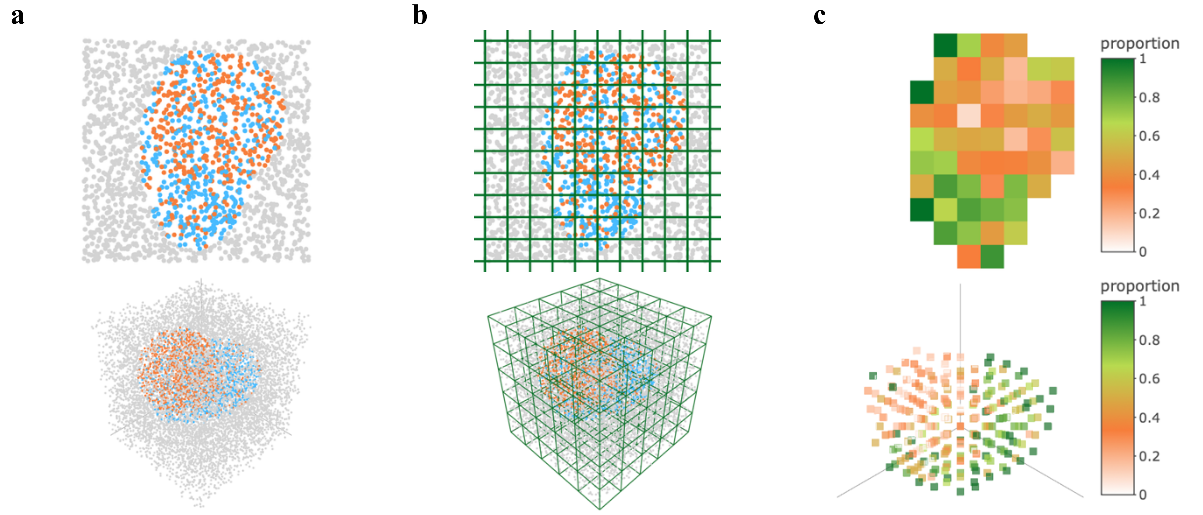


**Figure N1.7. Visualisation of grid metrics. (a)** 2D (top row) and 3D (bottom row) simulation of tissue structures containing tumour cells (orange), immune cells (blue) and background cells (grey). Generated using spaSim and spaSim-3D respectively. **(b)** Each simulation is broken down into a grid. The 2D simulation becomes composed of many rectangles while the 3D simulation becomes composed of many rectangular prisms. Each rectangle or rectangular prism can be individually examined. **(c)** Proportion grid metrics. Tumour cells were chosen as the reference cell type and immune cells were chosen as the target cell type and the proportion value in each grid rectangle/prism was calculated.

##### 1.2.1. Calculating metrics in each rectangular prism

Each rectangular prism in the 3D grid of the tissue will have a mixture of cells. Using the number of different cell types found in each rectangular prism, an individual metric value can be calculated for each rectangular prism to allow for further downstream spatial heterogeneity analysis.

If a set of reference cell types and target cell types are chosen by the user, then we can find the proportion of target cell types relative to reference and target cell types in each rectangular prism.

(15) $$\boldsymbol{Proportion\ of\ target\ cells\ in\ the\ xth\ grid\ prism} = \frac{\boldsymbol{n^x_{target}}}{\boldsymbol{n^x_{reference}} + \boldsymbol{n^x_{target}}}$$

Where:

- $n^x_{reference}$ is the number of reference cells in the $x$th rectangular prism.
- $n^x_{target}$ is the number of target cells in the $x$th rectangular prism.

On the other hand, if a single set of cell types of interest are chosen by the user, then the entropy of each rectangular prism can be calculated and visualised.

(16) $$\boldsymbol{P^x_i} = \frac{\boldsymbol{n^x_i}}{\sum_{i=1}^{n} \boldsymbol{n^x_i}}$$

(17) $$\boldsymbol{Entropy\ in\ the\ xth\ grid\ prism} = -\frac{\boldsymbol{1}}{\boldsymbol{\log_2(n)}} \sum_{i=1}^{n} \boldsymbol{P^x_i} \times \boldsymbol{\log_2(P^x_i)}$$

Where:

- $n_i^x$ is the number of $i$th cells in the $x$th rectangular prism.
- $P_i^x$ is the proportion of the $i$th cell in the $x$th rectangular prism.
- $n$ is the total number of cell types of interest.

Both metrics are useful in examining how cells are distributed throughout a tissue with the proportion value providing information about the relative amounts of reference and target cell types and the entropy value showing the levels of disorder of different cell types throughout a tissue (Figure N1.7c).

1.2.2. Prevalence and prevalence gradient

As described above, for each rectangular prism, a proportion value or entropy value can be calculated. By method of how these values are calculated, these values always range from 0 to 1. Therefore, a threshold between 0 and 1 can be specified by the user and the percentage of rectangular prisms with a proportion/entropy value above this threshold, or in other words, the 'prevalence' of a particular pattern can be calculated.[1]

However, similar to the gradient-based metrics implemented for several cell colocalization metrics, the prevalence can be graphed against a gradient of thresholds increasing from 0 to 1. Through this spatial heterogeneity metric, we can observe how pervasive particular spatial patterns are throughout a tissue.

1.2.3. Spatial autocorrelation

Using the metrics calculated for each rectangular prism as well as its spatial coordinates, spatial autocorrelation describes the degree to which proportion/entropy values at nearby locations are correlated to each other (i.e. how similar nearby prisms are). One way to calculate spatial autocorrelation is through Global Moran's I.[16]

(18) $$\boldsymbol{I} = \frac{\boldsymbol{n}}{\boldsymbol{W}} \frac{\sum_{i=1}^{n}\sum_{j=1}^{n} w_{ij}(x_i-\bar{x})(x_j-\bar{x})}{\sum_{i=1}^{n}(x_i-\bar{x})^2}$$

(19) $$\boldsymbol{W} = \sum_{i=1}^{n}\sum_{j=1}^{n} \boldsymbol{w_{ij}}$$

Where:

- $x_i$ and $x_j$ are the proportion/entropy value at the $i$th and $j$th rectangular prisms respectively.
- $\bar{x}$ is the mean proportion/entropy value across all rectangular prisms.
- $w_{ij}$ is the spatial weight matrix between the $i$th and $j$th rectangular prism.
- $n$ is the total number of rectangular prisms.

One of the inputs for the spatial autocorrelation formula is a spatial weight matrix, $w_{ij}$. The matrix defines which locations are considered 'neighbours' and how strongly they influence each other. Each cell in the matrix, $w_{ij}$, represents the spatial relationship between rectangular prism i and rectangular prism j. Typically, $w_{ij} = 1$ if rectangular prism i and j are neighbours, and $w_{ij} = 0$ otherwise. SPIAT-3D offers a range of different spatial weight matrices to choose from. For our analysis, we have chosen to use the 'queen' spatial weight matrix, where rectangular prisms are defined as neighbours if they share a common edge or corner.

Interpretation of Global Moran’s I is as follows:

1. Positive Global Moran’s I occurs when proportion/entropy values at nearby locations are similar, such as when rectangular prisms with high proportion values cluster together, and grid prims with low proportion values cluster together.
2. Negative Global Moran’s I when proportion/entropy values at nearby locations are dissimilar, such as when rectangular prisms with high entropy values surround themselves with rectangular prisms with low entropy values.
3. Zero Global Moran’s I suggests no link between proportion/entropy values at nearby locations and that the location of each rectangular prism is independent of their metric value.

In the context of spatial tissue data, Global Moran’s I provides a value for the overall spatial distribution of the rectangular prisms, revealing the level of clustering of cellular patterns throughout a tissue.

### Supplementary Note 2
### spaSim-3D

#### 2.1. Generating background cells

spaSim-3D uses a similar procedure to generate simulations as spaSim, but rather than just x and y coordinates, it also includes the z-coordinate of cells.[1]

To generate spatial patterns, spaSim first simulates 'background cells'. These are cells which are found throughout the simulated tissue space and which have not yet been given a cell type. Then, spaSim uses a Poisson point process for random generation of background cells. We have adapted this method for spaSim-3D by incorporating the z-coordinate of cells, enabling the simulation of background cells in 3D space.

##### 2.1.1. Algorithm for a Poisson point process in 3D to simulate a random pattern of background cells

To simulate background cells in a random pattern, spaSim employs a Poisson point process. In such distributions, each location contains a similar number of cells, and the positions of individual cells are independent.[17] Previous studies have shown that the position of cells exhibit a distribution following a Poisson point process in 2D cancer tissue sections.[1] To implement this, spaSim used the '*rHardcore*' function from the spatstat package[18] which can simulate a Poisson point process for a 2D plane. However, to our knowledge, there is no available package to implement this in 3D, and so we developed a novel algorithm for spaSim-3D which simulates the background cells in a 3D space following a Poisson point process.

The simulator requires the user to specify the desired number of cells and dimensions of the tissue space to be simulated. As we have set the tissue space to be a rectangular prism, the length, width and height of the tissue are required. Lambda (λ), a crucial parameter for the Poisson distribution, represents the average number of cells per unit volume and is set to five for spaSim-3D. While the choice of λ can impact the detailed characteristics of the cell distribution, our extensive testing suggests that it does not substantially influence the key features of interest in our simulations. Next, the 3D tissue is divided into a grid by splitting the x-axis, y-axis and z-axis into equal parts, forming a specific number of equally sized 3D rectangular prisms such that if every rectangular prism contained λ cells on average, the entire tissue would contain the number of cells requested by the user.

Calculations for the number of rectangular prisms and the corresponding number of rows, columns and layers in the grid needed are shown:

$$(20)\quad \boldsymbol{n_{prisms}} = \frac{\boldsymbol{n_{cells}}}{\boldsymbol{\lambda}} = \boldsymbol{n_{rows}} \times \boldsymbol{n_{cols}} \times \boldsymbol{n_{lays}}$$

$$(21)\quad \boldsymbol{n_{rows}} = \boldsymbol{n_{cols}} = \boldsymbol{n_{lays}} = \sqrt[3]{\boldsymbol{n_{prisms}}} = \sqrt[3]{\frac{\boldsymbol{n_{cells}}}{\boldsymbol{\lambda}}}$$

Where:

- $n_{rows}$, $n_{cols}$ and $n_{lays}$ are the number of rows, columns and layers used to divide the tissue along the x, y and z axis respectively into a 3D grid of rectangular prisms. For simplicity of this algorithm, we assume $n_{rows} = n_{cols} = n_{lays}$.
- $n_{prisms}$ is the total number of rectangular prisms in the grid.
- $n_{cells}$ is the number of cells in the tissue.
- $\lambda$ is the average number of cells per rectangular prism.

For each rectangular prism, a random number from a Poisson distribution with $\lambda = 5$ is generated, and that many cells are placed in that prism. By splitting the tissue into the grid following the above steps, the tissue is filled with the roughly the number of cells requested by the user, following a Poisson point process (Figure N2.1).

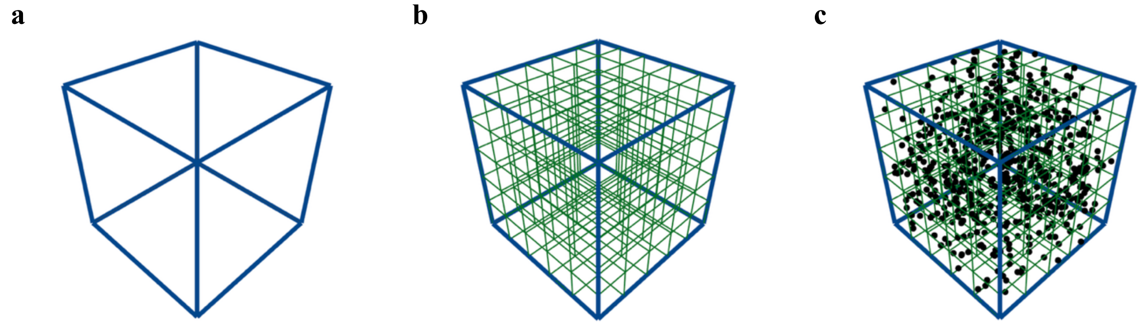


**Figure N2.1. Poisson point process to generate background cells in a 3D tissue. (a)** The rectangular prism tissue. **(b)** The tissue is divided into a 3D grid with a specific number of rectangular prisms. **(c)** Cells are placed randomly in each rectangular prism. The number of cells in each rectangular prism is obtained through a Poisson point process with $\boldsymbol{\lambda} = \mathbf{5}$.

Since the absolute smallest distance between each cell is limited by the size of the cell, another user parameter is called the 'minimum distance between cells'. For each cell whose distance to another cell is less than this chosen value, it is removed from the simulation. An example simulation of a random pattern background cells is shown in Figure N2.2a.

For our generated simulations, we wanted to make them as realistic as possible to existing biological tissue data. As the 'minimum distance between cells' parameter is a measure of cell size, this parameter needed to be chosen appropriately. Cells can range from 1μm to hundreds of micrometers. Based on our experience, we considered that 10μm was a reasonable average size for a cell. Therefore, we have set the 'minimum distance between cells' parameter to equal 10 throughout this work.

##### 2.1.2. Mixing the background cell composition

Up to now, background cells generated have no cell type identity assigned to them. If users wish to assign cell types randomly to the background cells, they can use the 'mixing' function. Here, the process is analogous to spaSim where the user inputs a selection of cell types and their desired proportions. Subsequently, each individual cell is reassigned to these chosen cell types with a probability that directly corresponds to the desired proportion of that cell type. An example of a mixed background is shown in Figure N2.2b.

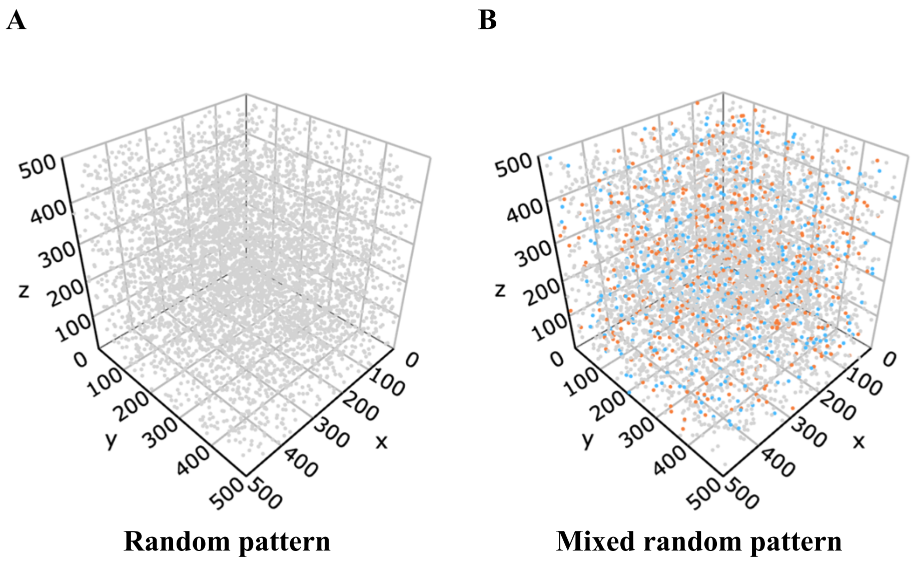


**Figure N2.2. Example background cell simulations.** The tissue size is 500 by 500 by 500, and the number of cells is 5,000 for both simulations. **(a)** Random pattern of background cells where the minimum distance between cells is 10. **(b)** The same random pattern in A but with mixing with 10% tumour cells (orange), 10% immune cells (blue) and 80% other cells.

#### 2.2. <u>Simulating differently shaped cell clusters</u>

spaSim-3D allows the user to generate cell clusters of a specified cell type by assigning a single cell type to regions of the previously simulated background cells. While many of these algorithms which simulate clusters have been extended from spaSim, novel methods for more intricate-shaped clusters have also been introduced to spaSim-3D. These clusters can differ in shape, size, and complexity, with the user having a variety of options to fully customise the clusters they choose to generate.

##### 2.2.1. <u>Sphere</u>

Similar to spaSim, which utilises a 2D Cartesian coordinate system $(x, y)$ to define circular clusters, our simulator employs a 3D Cartesian coordinate system $(x, y, z)$ to define spherical clusters within a 3D tissue background. The interior of the sphere is defined by the inequality:

(22) $$\boldsymbol{(x-a)^2 + (y-b)^2 + (z-c)^2 \leq r^2}$$

Where:

- $(x, y, z)$ are the coordinates of any point in space.
- $(a, b, c)$ are the coordinates of the sphere's centre.
- $r$ it the radius of the sphere.

To generate a sphere cluster in the simulation, the user first specifies the centre coordinate and radius of the sphere. Then, we take the $x$, $y$ and $z$ coordinate of each background cell and check if it satisfies the inequality for the interior of the sphere. For cells which do satisfy this inequality, the cell type identity of these cells can be updated to a particular cell type, such as a tumour cell. By customising the centre and radius of the sphere, the user has full control over the size and location of this sphere cluster in the tissue. An example is shown in Figure N2.3.

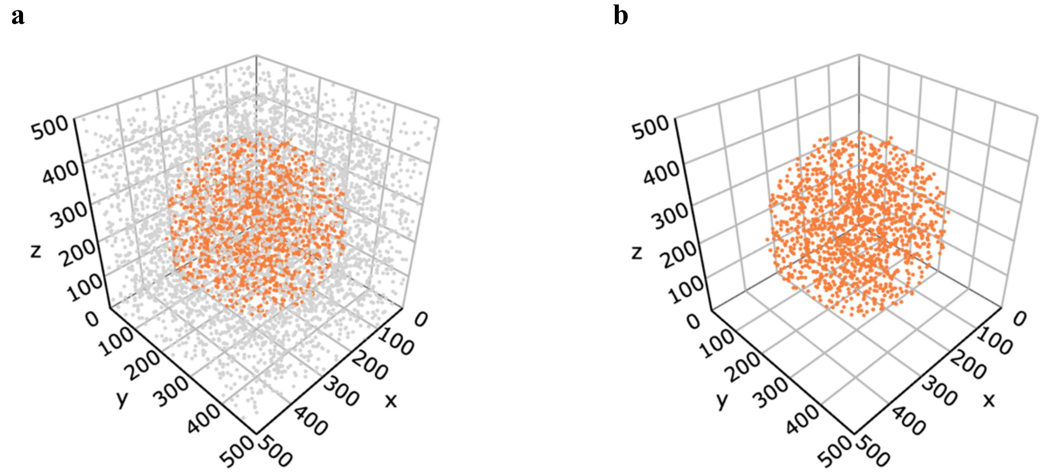


**Figure N2.3. Example sphere simulation.** A simulated tumour cell sphere where the radius is 200 and centre is (250, 250, 250). **(a)** Background cells simulated using a random pattern are included. The tissue size is 500 by 500 by 500, the number of cells is 5,000, and the minimum distance between cells is 10. **(b)** Background cells excluded.

##### 2.2.2. Ellipsoid

Building on the concept of a sphere, spaSim-3D can also simulate ellipsoidal clusters and allows users to control their orientation. While spaSim previously simulated ellipses in a 2D Cartesian coordinate system, we have extended this approach to 3D, allowing for the generation of ellipsoidal clusters in a 3D tissue background. The interior of the ellipsoid is defined by the inequality:

$$(23) \quad \frac{(x-a)^2}{r_x^2} + \frac{(y-b)^2}{r_y^2} + \frac{(z-c)^2}{r_z^2} \leq 1$$

Where:

- $(x, y, z)$ are the coordinates of any point in space.
- $(a, b, c)$ are the coordinates of the ellipsoid's centre.
- $r_x$ is the semi-axis length of the ellipsoid in the *x* direction.
- $r_y$ is the semi-axis length of the ellipsoid in the *y* direction.
- $r_z$ is the semi-axis length of the ellipsoid in the *z* direction.

To generate an ellipsoidal cluster, the user first specifies the centre coordinate and semi-axes lengths of the ellipsoid. Then spaSim-3D changes the cell type identity of background cells whose coordinates satisfy the inequality defining the interior of the ellipsoid.

However, spaSim's implementation of elliptical clusters and correspondingly, the current ellipsoid inequality, assumes a fixed orientation aligned with the coordinate axes. To introduce greater flexibility and mimic the varying orientations observed in biological structures, the user can also choose the ellipsoid's angle of rotation in the *y*-*z*, *x*-*z*, and *x*-*z* planes. Due to the introduction of ellipsoid rotation, we could no longer use the above ellipsoid inequality and instead developed a new method involving matrix transformations to simulate a fully transformed ellipsoid. This process is shown in Figure N2.4.

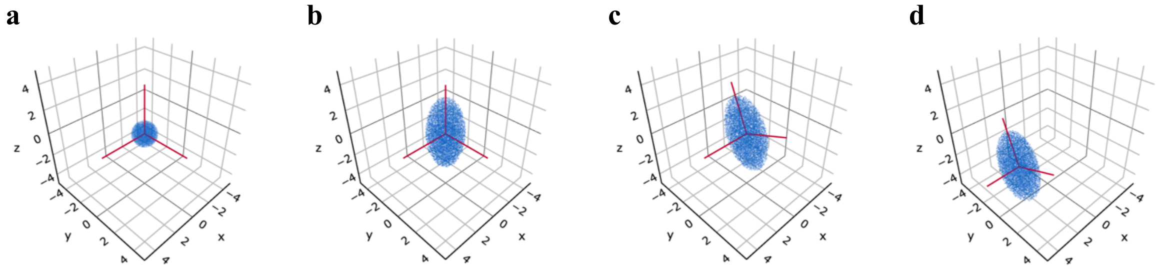

**Figure N2.4. Conversion of a unit sphere to a fully transformed ellipsoid. (a)** Unit sphere centred at the origin. **(b)** Unit sphere after dilation along each axis. **(c)** Unit sphere after dilation and rotation. **(d)** Unit sphere after dilation, rotation and translation to become a fully transformed ellipsoid.

The unit sphere is first dilated to a simple ellipsoid. The dilation matrix for a 3D ellipsoid with semi-axes lengths of $r_x$ , $r_y$ , and $r_z$ is:

$$(24) \quad \boldsymbol{T_0} = \begin{bmatrix} \boldsymbol{r_x} & \boldsymbol{0} & \boldsymbol{0} \\ \boldsymbol{0} & \boldsymbol{r_y} & \boldsymbol{0} \\ \boldsymbol{0} & \boldsymbol{0} & \boldsymbol{r_z} \end{bmatrix}$$

Then, the ellipsoid is rotated along each plane. The transformation matrices for rotation in the *y-z* plane ($T_1$) by theta ($\theta$), *x-z* plane ($T_2$) by alpha ($\alpha$) and *x-y* plane ($T_3$) by beta ($\beta$) are shown:

$$(25) \quad \boldsymbol{T_1} = \begin{bmatrix} \boldsymbol{1} & \boldsymbol{0} & \boldsymbol{0} \\ \boldsymbol{0} & \boldsymbol{\cos(\theta)} & \boldsymbol{-\sin(\theta)} \\ \boldsymbol{0} & \boldsymbol{\sin(\theta)} & \boldsymbol{\cos(\theta)} \end{bmatrix}$$

$$(26) \quad \boldsymbol{T_2} = \begin{bmatrix} \boldsymbol{\cos(\alpha)} & \boldsymbol{0} & \boldsymbol{-\sin(\alpha)} \\ \boldsymbol{0} & \boldsymbol{1} & \boldsymbol{0} \\ \boldsymbol{\sin(\alpha)} & \boldsymbol{0} & \boldsymbol{\cos(\alpha)} \end{bmatrix}$$

$$(27) \quad \boldsymbol{T_3} = \begin{bmatrix} \boldsymbol{\cos(\beta)} & \boldsymbol{-\sin(\beta)} & \boldsymbol{0} \\ \boldsymbol{\sin(\beta)} & \boldsymbol{\cos(\beta)} & \boldsymbol{0} \\ \boldsymbol{0} & \boldsymbol{0} & \boldsymbol{1} \end{bmatrix}$$

Finally, the ellipsoid is translated. The translation needed to get an ellipsoid of centre at $(a, b, c)$ is acquired by adding the following matrix to a point $(x, y, z)$:

$$(28) \quad \boldsymbol{T_4} = \begin{bmatrix} \boldsymbol{a} \\ \boldsymbol{b} \\ \boldsymbol{c} \end{bmatrix}$$

Applying these matrices to a point $(x, y, z)$ yields the coordinates of the transformed point $(x_{new}, y_{new}, z_{new})$ and the following equation:

$$(29) \quad \boldsymbol{T_3 T_2 T_1 T_0} \begin{bmatrix} \boldsymbol{x} \\ \boldsymbol{y} \\ \boldsymbol{z} \end{bmatrix} + \boldsymbol{T_4} = \begin{bmatrix} \boldsymbol{x_{new}} \\ \boldsymbol{y_{new}} \\ \boldsymbol{z_{new}} \end{bmatrix}$$

Solving for $(x, y, z)$ yields:

$$(30) \quad \begin{bmatrix} \boldsymbol{x} \\ \boldsymbol{y} \\ \boldsymbol{z} \end{bmatrix} = \boldsymbol{T_0^{-1} T_1^{-1} T_2^{-1} T_3^{-1}} \left( \begin{bmatrix} \boldsymbol{x_{new}} \\ \boldsymbol{y_{new}} \\ \boldsymbol{z_{new}} \end{bmatrix} - \boldsymbol{T_4} \right)$$

Hence, equations for $x$, $y$, and $z$ in terms of $x_{new}$, $y_{new}$, and $z_{new}$ can be obtained and substituted into the inequality for the interior of a unit sphere:

(31) $$\boldsymbol{x^2 + y^2 + z^2 \leq 1}$$

This yields the inequality representing a fully transformed ellipsoid.

By checking if any cell in the background has coordinates which satisfy this transformed inequality, the user is able to simulate an ellipsoid cluster with complete control over its size, position and orientation. An example is shown in Figure N2.5.

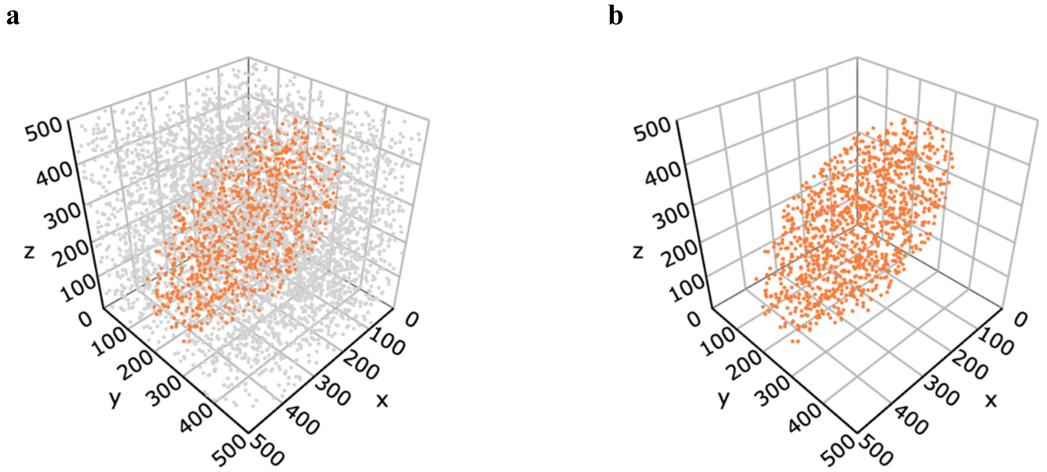


**Figure N2.5. Example ellipsoid simulation.** A tumour ellipsoid where semi-axes lengths are 150, 150 and 300, centre is (250, 250, 250) and x-z plane rotation is 60º. **(a)** Background cells simulated using a random pattern. The tissue size is 500 by 500 by 500, the number of cells is 5,000, and the minimum distance between cells is 10. **(b)** Background cells excluded.

2.2.3. <u>Cylinder</u>

To simulate vessels and tumour areas that have grown in shapes resembling cylinders, the user can simulate cylinders, and can customise its shape by inputting the desired thickness or radius, start position and end position in spaSim-3D.

Conceptually, a cell is inside the cylinder if it satisfies two conditions (Figure N2.6):

2. The perpendicular distance of the cell to the axis of the cylinder (i.e. the line that passes through both the start and end location) is less than the cylinder's radius.
3. A cell is contained between the planes drawn at the start and end locations of the cylinder and perpendicular to the axis of the cylinder.

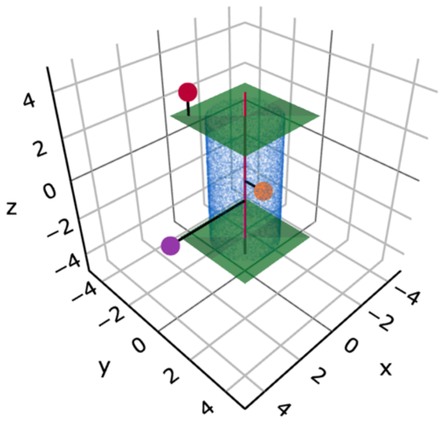


**Figure N2.6**. **Determining if a cell is inside the cylinder.** The red line represents the cylinder's axis. The purple cell is outside the cylinder as its perpendicular distance to the cylinder's axis is greater than the cylinder's radius. The red cell is also outside the cylinder as it is on the outside of the top plane that caps the cylinder. The orange cell is inside the cylinder as it satisfies both conditions.

To accomplish this mathematically, we can use a vector algebra formulae derived from linear algebra equations.[19]

Let $OP = (x, y, z)$ represent any point in space, $OA = (x_0, y_0, z_0)$ the coordinates of the cylinder's start location, $OB = (x_1, y_1, z_1)$ the coordinates of the cylinder's end location and $r$ its radius.

The vector between OA and OB is given by:

(32) $\boldsymbol{AB} = (\boldsymbol{x_1} - \boldsymbol{x_0}, \boldsymbol{y_1} - \boldsymbol{y_0}, \boldsymbol{z_1} - \boldsymbol{z_0})$

The vector between OP and OA is given by:

(33) $\boldsymbol{AP} = (\boldsymbol{x} - \boldsymbol{x_0}, \boldsymbol{y} - \boldsymbol{y_0}, \boldsymbol{z} - \boldsymbol{z_0})$

The perpendicular distance, d, from P to the line between A and B is given by:

(34) $\boldsymbol{d} = \frac{|\boldsymbol{AP} \times \boldsymbol{AB}|}{|\boldsymbol{AB}|}$

Therefore, the first condition for a cell to be inside the cylinder, that the perpendicular distance of the cell to the cylinder's axis is less than the cylinder's radius, can be represented by the following inequality:

(35) $\boldsymbol{d} \leq \boldsymbol{r}$

Also, the vector equations for the planes at A and B are given by:

(36) $\boldsymbol{OP} \cdot \boldsymbol{AB} = \boldsymbol{OA} \cdot \boldsymbol{AB}$

(37) $\boldsymbol{OP} \cdot \boldsymbol{AB} = \boldsymbol{OB} \cdot \boldsymbol{AB}$

Therefore, second first condition for a cell to be inside the cylinder, that the cell is located between the two planes at both ends of the cylinder, can be represented by the following inequality:

(38) $\boldsymbol{OA} \cdot \boldsymbol{AB} \leq \boldsymbol{OP} \cdot \boldsymbol{AB} \leq \boldsymbol{OB} \cdot \boldsymbol{AB}$

By combining cylinders with other shapes, complex 3D tissues can be simulated, containing both tumour masses and blood vessels structures. An example is shown in Figure N2.7.

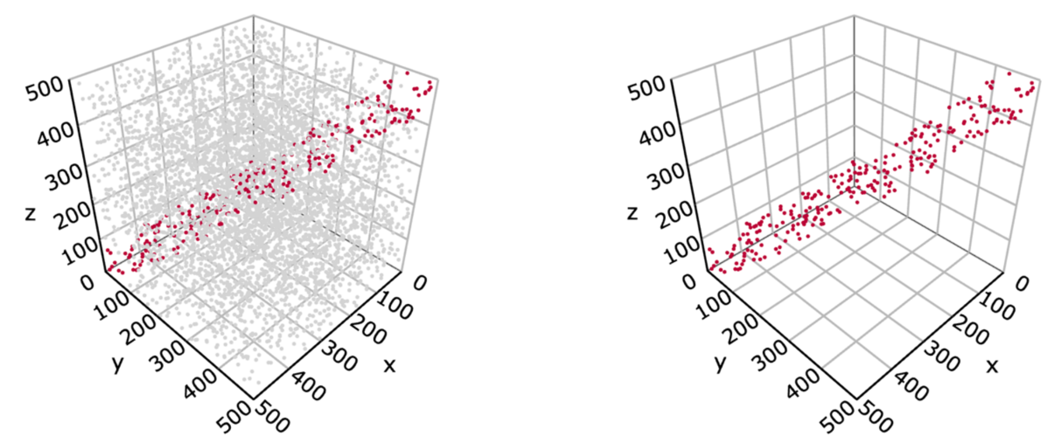


**Figure N2.7**. **Example cylinder simulation.** A cylinder blood vessel where radius is 50, start point is (500, 0, 0) and end point is (0, 500, 500). **(a)** Background cells simulated using random pattern. The tissue size is 500 by 500 by 500, the number of cells is 5,000, and the minimum distance between cells is 10. **(b)** Background cells excluded.

2.2.4. Network-like structure

Despite the large amount of diversity in spatial patterns that can be generated from using spheres, ellipsoids and cylinders, they still fall short of simulating more intricate and irregular structures seen in 3D tissues. For instance, Lin et al.[8] showcased the formation of tertiary lymphoid structure networks in a 3D colorectal cancer sample. Therefore, we designed spaSim-3D to also be able to simulate 3D network-like structures. This is a novel feature that was not included in the original spaSim implementation.

We first assumed that a network-like structure is comprised of different vertices all connected to each other by edges, and located in a fixed space in the tissue. The parameters needed by the user include the number of edges in the network, the width of each edge, the centre of the network-like structure, and the radius of the sphere in which it is contained (containment radius).

To determine how the vertices of the network would be connected, we further assumed the linkages within the network would follow a minimum spanning tree (MST).[7]

If we start off with every single vertex being connected to every single other vertex with an edge, an MST only uses a specific subset of those edges. Furthermore, in MST, each edge must have a numeric value, or 'weight' assigned to it (e.g. cost, distance). For an MST, the edges are chosen such that:

1. There are no cycles between vertices (cannot start at a vertex and return to it by following distinct edges).

2. All vertices are still connected to each other (a path exists between every pair of vertices, so no vertex is isolated).
3. The sum of the edge weights is a minimum.

Due to the requirements of an MST, the number of vertices is always one more than the number of edges in an MST. Therefore, this many coordinates are randomly chosen from inside the sphere that the network is contained in to represent the vertices of the MST (Figure N2.8).

For the weight of each edge, we chose this to be the Euclidean distance between the vertices.

Finally, to construct the MST, we employed Prim's algorithm.[7] The algorithm starts off by choosing an arbitrary starting vertex to form the basis of the MST. Then, it finds the vertex that is closest to this starting vertex (the edge with the smallest weight) and adds it to the MST. From all connected vertices that are part of the MST, the algorithm adds the next closest, unconnected vertex to the MST and repeats this process until all vertices are connected. Prim's algorithm efficiently identifies the subset of edges that form the MST (Figure N2.8).

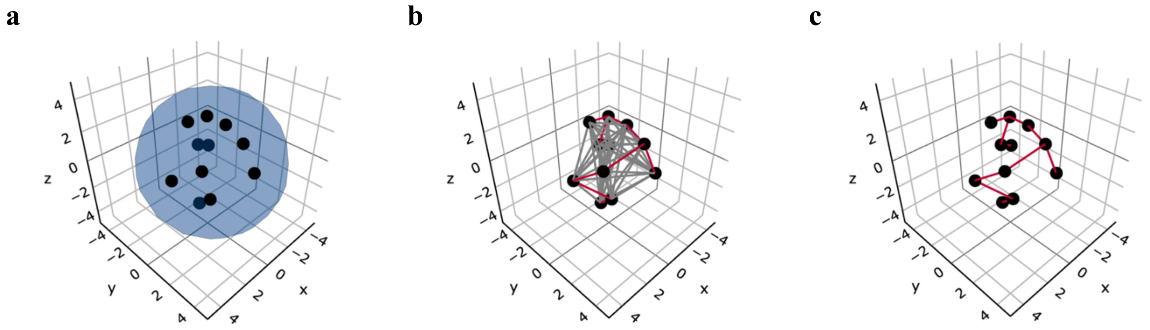


**Figure N2.8. Construction of an MST in 3D space. (a)** Random points inside the chosen sphere are chosen to represent the MST vertices. **(b)** Every vertex is connected to every other vertex with an edge. The MST of the graph is shown in red. **(c)** The subset of the edges showing only the edges that form the MST.

With the constructed MST, we can simulate the network-like structure. To do this, a cylinder is simulated for each edge. The previous cylinder simulator function (Supplementary Note 2, Section 2.2.3) requires a start position, end position and thickness. Here, the coordinates of the vertices connected by one of the edges are used as the start and end coordinates of the cylinder, and the user-inputted edge width parameter is used as the thickness of the cylinder. By simulating a cylinder for each edge, a network-like structure can be generated (Figure N2.9).

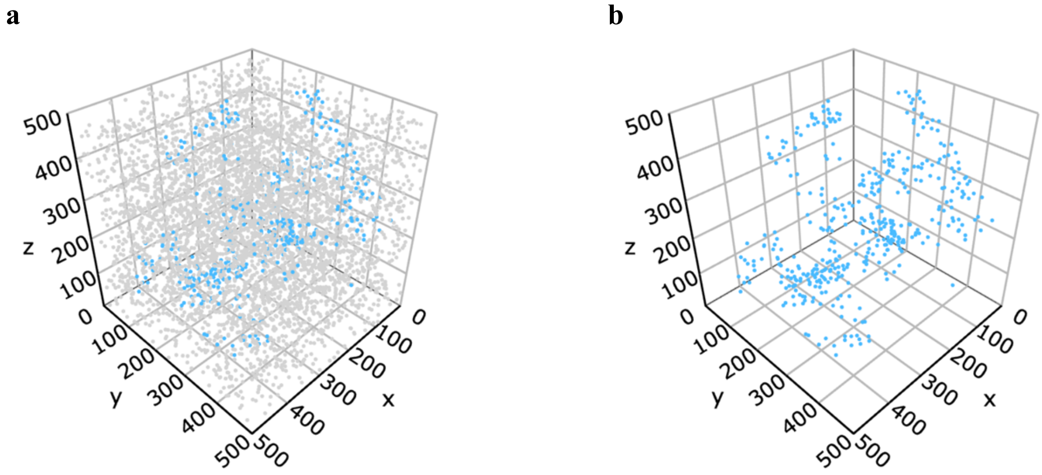


**Figure N2.9. Example network simulation.** Network-like structure where the containment radius is 300, centre is (250, 250, 250), number of edges is 50, width of each edge is 40. **(a)** Background cells simulated using a random pattern. The tissue size is 500 by 500 by 500, the number of cells is 5,000, and the minimum distance between cells is 10. **(b)** Background cells excluded.

#### 2.3. Customising clusters

While the selection of different 3D shapes and structures gives the user the ability to generate a wide variety of 3D spatial patterns, we included additional features to further customise the complexity of the clusters to better reflect what is observed in biological tissue structures.

##### 2.3.1. Mixing the cluster cell composition

The clusters depicted previously consisted of a single cell type, such as only tumour cells or only immune cells. However, real tissue structures often comprise a diverse mix of cell types in varying proportions, as exemplified by tumour masses infiltrated by immune cells. To implement this in spaSim-3D, users can input a selection of cell types and their desired proportions for the cluster. Subsequently, the cells within the cluster are reassigned to these chosen cell types with a probability that directly corresponds to the desired proportion of each cell type, as illustrated in Figure N2.10a.

##### 2.3.2. Adding rings to clusters

Another common cell arrangement observed in tissues is the formation of cell rings at the periphery of clusters. For example, in the phenomenon known as immune exclusion, immune cells are recruited around the perimeter of tumour clusters.[20] Given that this feature was already present in spaSim, we have directly implemented spaSim's procedure for simulating cell rings within spaSim-3D. spaSim also facilitates simulation of double rings around clusters which gave the user the ability to vary the composition and arrangement of cells at the outer-edge of clusters – this feature has also been transferred to spaSim-3D.

Along with specifying the parameters for a cluster, such as its shape, size and position in the tissue, the user can also specify ring-specific parameters. These include the width of the ring as well as the different cell types and corresponding cell proportions that comprise the ring. To simulate a cluster with a ring, spaSim-3D first simulates a larger cluster which takes into account the ring width (i.e. the ring width is added to the radius of the sphere, ellipsoid or cylinder, or is added to the width of a network branch when simulating these clusters). Then, a smaller cluster without the ring width is simulated inside of this larger cluster, giving the illusion of a ring. To simulate clusters with a double ring in spaSim-3D, the user provides ring-specific parameters for both an outer ring and inner ring. Then, a cluster which uses both the inner and outer ring is simulated first, a cluster with only the inner ring is simulated next, and finally a cluster without either of these rings is simulated, creating a double ring. Examples are shown in Figure N2.10b and Figure N2.10c.

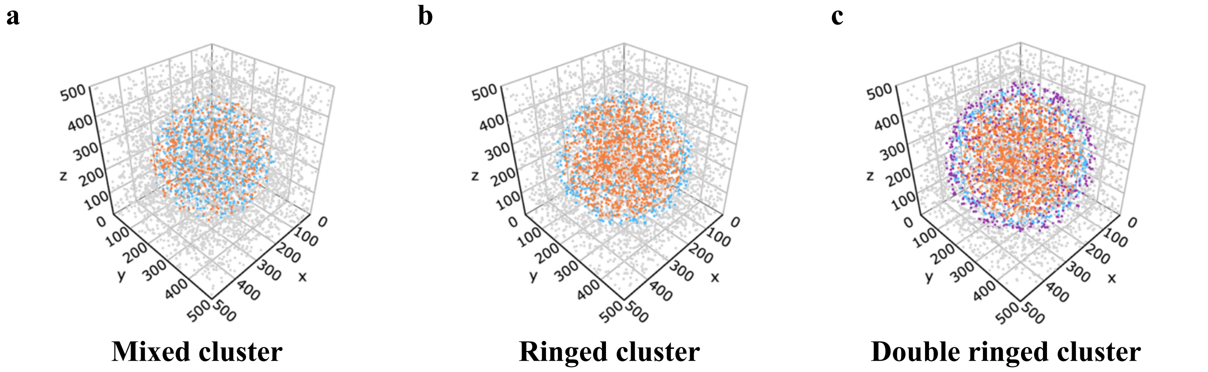


**Figure N2.10**. **Example cluster simulations with further customisation.** For each simulation, the same random pattern was used to simulate the background cells. The tissue size is 500 by 500 by 500, the number of cells is 5,000, and the minimum distance between cells is 10. The same sphere cluster was also generated for each where the radius is 175 and centre is (250, 250, 250). **(a)** Cluster is mixed with 50% tumour cells (orange) and 50% immune cells (blue). **(b)** Tumour cluster (orange) with an immune ring (blue) of width 20. **(c)** Tumour cluster (orange) with an 'immune1' inner ring (blue) of width 20 and an 'immne2' outer ring (purple) of width 20.
