## Supplementary Tables for "2D spatial tissue analysis often misrepresents true biological patterns of 3D tissues"

**Table S1 legend. Summary values of median percentage differences between 3D and 2D spatial metrics in publicly available 3D datasets.** The percentage difference was calculated as: $\boldsymbol{percentage\ difference} = \left(\frac{\boldsymbol{2D\ value - 3D\ value}}{\boldsymbol{3D\ value}}\right) \times \mathbf{100\%}$. 3D metrics were applied to each 3D dataset while 2D equivalent metrics were applied to each extracted slice from the same 3D dataset. Multiple cell pairs (e.g. Tumour/Tumour, Tumour/Immune, etc.) were used as inputs for the 3D and 2D metrics. For each slice, the percentage difference between the 3D and 2D metric values was computed for every cell pair. The median percentage difference across all cell pairs was then taken for that slice. The minimum, maximum, range, median and average of these slice-based medians are shown for human colorectal cancer sample generated with CyCIF by Lin et al.,[8] human metastatic lymph node sample generated with Open-ST by Schott et al.[9] and mouse brain cortex and hypothalamus generated with MERFISH by Fang et al.[10]

**Table S2 legend. Spearman correlations between 3D metric values and 2D metric values for Simulated Dataset 1.** 3D metric values were obtained by applying each 3D metric to the original 3D simulation. For 'random slice', 2D metric values were obtained by extracting 13 slices from each simulation and applying each 2D metric to a randomly sampled slice. For 'average slice', 2D metric values were obtained by applying each 2D metric to all 13 slices from each simulation and then taking the average. For 'middle slice', 'upper slice' and 'uppermost slice', 2D metric values were obtained by applying each 2D metric to the middle slice (z = 145 to z = 155), upper slice (z = 175 to z = 185) and uppermost slice (z = 205 to z = 215) respectively. For 'mixed-ellipsoid', 'mixed-network', 'ringed-ellipsoid', 'ringed-network', 'separated-ellipsoid', 'separated-network', 2D metric values were obtained by first subsetting the simulations for the corresponding tissues structure (e.g. mixed-ellipsoid, 'mixed-network', etc.), then applying each 2D metric to a randomly sample slice from each simulation subset.

**Table S3 legend. Summary values of percentage differences between 3D and 2D metrics for Simulated Dataset 1.** The percentage difference was calculated as: $\boldsymbol{percentage\ difference} = \left(\frac{\boldsymbol{2D\ value - 3D\ value}}{\boldsymbol{3D\ value}}\right) \times \mathbf{100\%}$. Data collected for 3D metric values and 2D metric values is the same as the data collected for 'random slice' and 'average slice' in Table S2. The minimum, maximum, range, median and average percentage difference values are shown.

**Table S4. Fixed parameters used in spaSim-3D simulations.**

| Simulation feature | Parameter | Description | Value(s) |
|---|---|---|---|
| **Background** | dimensions | Length, width and height of the background simulation space | 600 by 600 by 300 |
| | n_cells | Number of cells | 30,000 |
| | min_distance | Minimum distance between cells | 10 |
| **Arrangement specific features** | cluster_centre | Centre coordinate of clusters | (300, 300, 150)[a] |
| | cluster2 | For separated clusters: Cluster 2 is a fixed sphere | Radius: 100 Centre: (450, 300, 150) |
| **Shape specific features** | N_n_edges | For networks: Number of edges | 20 |
| | N_radius | For networks: Radius of containment sphere | 125 |

[a]For separated clusters, the x-coord of cluster[1] is not equal to the fixed value of 300 but varies according to the cluster1_x_coord parameter in **Table S6**.

**Table S5. Categorical parameters used in spaSim-3D simulations.**

| Simulation feature | Parameter | Description | Value(s) |
|---|---|---|---|
| **Cluster** | arrangement | Arrangement of cells in cluster | Mixed, ringed, or separated |
| | shape | Shape of cluster | Ellipsoid or network |

**Table S6. Continuous parameters used in spaSim-3D simulations.**

| Simulation feature | Parameter | Description | Fixed value[a] | Range |
|---|---|---|---|---|
| **Background** | bg_prop_A[c] | Proportion of cell type A in the background | 0 | (0, 0.1)[b] |
| | bg_prop_B[c] | Proportion of cell type B in the background | 0 | (0, 0.1) |
| **Arrangement specific features** | cluster_prop_A[d] | For mixed clusters: The proportion of cell type A in the cluster | 0.5 | (0.3, 0.7) |
| | ring_width_factor[e] | For ringed clusters: Multiplying factor to get the ring width | 0.15 | (0.1, 0.2) |
| | cluster1_x_coord | For separated clusters: x-coord of cluster 1 | 150 | (125, 175) |
| **Shape specific features** | E_radius_x[f] | For ellipsoids: Radius in x-axis | 75 | (75, 125) |
| | E_radius_y[f] | For ellipsoids: Radius in y-axis | 100 | (75, 125) |
| | E_radius_z[f] | For ellipsoids: Radius in z-axis | 125 | (75, 125) |
| | N_width | For networks: Width of edge | 30 | (25, 35) |

[a]Fixed values used for Simulated Dataset 2 are found by taking the middle value of the corresponding continuous interval, except for bg_prop_A and bg_prop_B, which have been set to zero to ensure no background noise when testing other parameters.
[b](a, b) represents a continuous interval between a and b. A random value was uniformly sampled from this interval for each parameter.
[c]The background of each simulation is composed of cell type O (the background cell type), as well as cell type A and B, depending on their proportion in the background. The proportion of cell type O in the background (bg_prop_O) can be calculated by 1 – bg_prop_A – bg_prop_B.
[d]For mixed clusters, the proportion of cell type B in the cluster (simulation parameter for spaSim-3D) can be calculated by 1 – cluster_prop_A.
[e]For ringed clusters, the ring width (simulation parameter for spaSim-3D) is calculated by multiplying the ring_width_factor by the average of the ellipsoid radii or the network width.
[f]E_radius_x, E_radius_y and E_radius_z were collectively considered as one continuous parameter. These 3 parameters were all varied simultaneously, or all kept fixed.

**Table S7. Continuous parameters changed within each categorical parameter set in Simulated Dataset Two**. Categorical parameter sets are derived from combinations of categorical parameters in Table S5. Description of each continuous parameter is shown in Table S6.

| Categorical parameter set | Background | Arrangement specific features | Shape specific features |
|---|---|---|---|
| **mixed-ellipsoid** | bg_prop_A & bg_prop_B | cluster_prop_A | E_volume[a] |
| **mixed-network** | bg_prop_A & bg_prop_B | cluster_prop_A | N_width |
| **ringed-ellipsoid** | bg_prop_A & bg_prop_B | ring_width_factor | E_volume |
| **ringed-network** | bg_prop_A & bg_prop_B | ring_width_factor | N_width |
| **separated-ellipsoid** | bg_prop_A & bg_prop_B | distance[b] | E_volume |
| **separated-network** | bg_prop_A & bg_prop_B | distance | N_width |

[a]For ellipsoid clusters, 'E_volume' refers to the volume of the ellipsoid. Its three radii values (Table S6) can be inputted into the formula for the volume of the ellipsoid to calculate its volume.
[b]For separated clusters, 'distance' refers to the distance between cluster1 and cluster2 and can be found by taking the difference between their x-coords (Table S6).

**Table S8. Spatial metrics applied to each simulation.**

Each metric requires an input of cell types, typically a reference and target cell type, which was chosen to be the A/B cell pair, where the reference cell type was set to cell type 'A' and the target cell type was set to cell type 'B'. Cell colocalization metrics which use an input gradient of radius values (ANC, ACIN, ANE, MS, NMS, CK, CL, CG, COO) are applied to each simulation for a gradient of radius values from 20 to 100 with a step size of 10. Then, the area under the curve is calculated across the gradient of radius values using the Trapezoidal Rule. Spatial heterogeneity metrics (PBSAC, PBP, EBSAC, EBP) require a number of splits to divide each 3D tissue sample, which was chosen to be 10 splits along each axis (x, y, and z). Spatial heterogeneity metrics which use an input gradient of threshold values (PBP, EBP) are applied to each simulation for a gradient of threshold values from 0.01 to 1 with a step size of 0.01. Then, the area under the curve is calculated across the gradient of threshold values using the Trapezoidal Rule.

| Metric | Cell colocalization | Requires gradient of radius values | Spatial heterogeneity | Requires gradient of threshold values |
|---|---|---|---|---|
| Average minimum distance (AMD) | ✓ | × | × | × |
| Average neighbourhood counts (ANC) | ✓ | ✓ | × | × |
| Average cells in neighbourhood (ACIN) | ✓ | ✓ | × | × |
| Average neighbourhood entropy (ANE) | ✓ | ✓ | × | × |
| Mixing score (MS) | ✓ | ✓ | × | × |
| Normalised mixing score (NMS) | ✓ | ✓ | × | × |
| Cross K function (CK) | ✓ | ✓ | × | × |
| Cross L function (CL) | ✓ | ✓ | × | × |
| Cross G function (CG) | ✓ | ✓ | × | × |
| Co-occurrence (COO) | ✓ | ✓ | × | × |
| Proportion-based spatial autocorrelation (PBSAC) | × | × | ✓ | × |
| Proportion-based prevalence (PBP) | × | × | ✓ | × |
| Entropy-based spatial autocorrelation (EBSAC)[a] | × | × | ✓ | ✓ |
| Entropy-based prevalence (EBP)[a] | × | × | ✓ | ✓ |

[a]The EBSAC and EBP metrics uses a 'cell types of interest' input, rather than a separate reference and target cell type input. The 'cell types of interest' input for EBSAC and EBP was set to both cell type A and B.

**Table S9. Parameter ranges used in spaSim-3D simulations for Simulated Dataset 3.**
In Simulated Dataset 3, each collection is composed of simulations with fixed parameter values from Table S4 and fixed values of continuous variable parameters from Table S6. The categorical parameters and ranges of variable parameter used for each group of each collection are shown here.

| Collection | FPRc[a] or FNRc[b] | Group 1 variable parameter | Group 1 variable parameter range | Group 2 variable parameter | Group 2 variable parameter range |
|---|---|---|---|---|---|
| Mixed ellipsoids vs mixed ellipsoids | FPRc | cluster_prop_A[c] | (0.3, 0.7) | cluster_prop_A | (0.3, 0.7) |
| Mixed networks vs mixed networks | FPRc | cluster_prop_A | (0.3, 0.7) | cluster_prop_A | (0.3, 0.7) |
| Mixed ellipsoids vs mixed ellipsoids | FNRc | cluster_prop_A | (0.3, 0.7) | cluster_prop_A | (0.5, 0.9) |
| Mixed networks vs mixed networks | FNRc | cluster_prop_A | (0.3, 0.7) | cluster_prop_A | (0.5, 0.9) |

[a]A false positive rate collection (FPRc) tests the FPR, as the parameter range used in group 1 and group 2 are the same.
[b]A false negative rate collection (FNRc) tests the FNR, as the parameter range used in group 1 and group 2 are different.
[c]cluster_prop_A refers to the proportion of cell type A in the mixed cluster. The proportion of cell type B in the cluster (simulation parameter for spaSim-3D) can be calculated by 1 – cluster_prop_A.

**Table S10 legend. Data points excluded from figures for Simulated Dataset 1 for simulations where all 13 slices were extracted and an average 2D metric value across the slices was obtained.** Data points from Supplementary Figure S5a, Supplementary Figure S6b which contained percentage difference values below -1000% or above 1000% are stated.

**Table S11 legend. Data points excluded from figures for Simulated Dataset 1 for simulations where 3 slices (middle, upper and uppermost) were extracted.** Data points from Supplementary Figure S5b, Supplementary Figure S6c which contained percentage difference values below -1000% or above 1000% are stated.

**Table S12 legend. Data points excluded from figures for Simulated Dataset 1 for simulations where a random slice was extracted.** Data points from Figure 1d, Supplementary Figure S5c, Supplementary Figure S6a and Supplementary Figure S6d which contained percentage difference values below -1000% or above 1000% are stated.
